# Staying in the loop to make ends meet: roles and regulation of GlmR in *Bacillus subtilis*

**DOI:** 10.1101/2025.11.05.686802

**Authors:** Logan Suits, Sebastian J. Khan, Dipanwita Bhattacharya, Silviya Dimitrova, Prahathees J. Eswara

## Abstract

The metabolic networks of most life forms integrate cost-benefit analysis to properly budget carbon and other essential nutrients through continuous assessment of nutrient availability and environmental threats. *Bacillus subtilis* is a Gram-positive model bacterium found in diverse ecological niches such as soil, marine environments, and the human gut. As such, *B. subtilis* cells finetune metabolic pathways by monitoring signals indicating the presence of nutrients and stressors. A highly conserved protein, GlmR, is a key player in rationing carbon for the production of cell envelope precursors. This function of GlmR can be attributed to its role in cell shape regulation and antibiotic resistance. Given its central position in carbon utilization, GlmR is under post-translational regulation by phosphorylation and UDP-N-acetylglucosamine (UDP-GlcNAc) binding. GlmR is also linked to cyclic-di-AMP (c-di-AMP), a nucleotide second messenger involved in stress response. In this study, we probed the importance of GlmR in cell morphogenesis, c-di-AMP signaling, and investigated the physiological significance of post-translational regulation. Our results reveal that cells lacking *glmR* exhibit: (i) increased susceptibility to tunicamycin, a cell envelope targeting antibiotic; (ii) impaired division site positioning; and (iii) elevated intracellular c-di-AMP concentration. Furthermore, we show that the function of GlmR is finetuned by UDP-GlcNAc binding, phosphorylation, and acetylation. Additionally, we provide evidence showing that the recently discovered enzymatic activity of GlmR is integral for its function. We show that GlmR is a cell width determinant and propose a model suggesting close cooperation with an actin-like protein, MreB. Overall, our studies highlight that GlmR is at the crux of carbon flux with an important role in maintaining cell envelope integrity.

**IMPORTANCE:** Bacteria must integrate feedback from multiple metabolic processes to efficiently allocate carbon to produce essential building blocks such as nucleotides, amino acids, and cell wall precursors to support life. GlmR is a critical metabolic factor involved in the making of cell envelope precursors in diverse bacterial phyla. In *Bacillus subtilis*, cells lacking GlmR are deformed and hypersensitive to cell wall targeting antibiotics. As siphoning off too much carbon from other essential processes is detrimental to cell viability, GlmR activity is tightly regulated. Here we report that absence of GlmR leads to aberrant placement of cytokinetic machinery and an increase in the levels of cyclic-di-AMP, a nucleotide second messenger that assists in the cell wall stress response. We also show that GlmR function is post-translationally finetuned by phosphorylation and acetylation. Furthermore, our data reveals that the catalytic activity of GlmR is required for its function. Thus, the activity of GlmR is tightly calibrated through multiple means for efficient carbon utilization.

## INTRODUCTION

To survive and thrive, bacteria must be adept at tuning the metabolic pathways to shuttle carbon for building genetic material, cell envelope, and generating energy. This regulation must happen within seconds to assess, calibrate, and shunt metabolic intermediates to appropriate biosynthetic cycles. Post-translational regulation of metabolic proteins and the use of second messengers greatly aid in achieving this instantaneous response. *Bacillus subtilis* is a common soil bacterium generally considered as the model for Gram-positive organisms (1). Many bacteria including *B. subtilis* occupy and adapt to diverse ecological niches including the human gut (2) and marine environments (3). Therefore, *B. subtilis* cells must skillfully tackle a multitude of challenges such as varying levels of nutrients, osmotic shifts, and other types of stressors. To achieve this, they must tightly ration carbon for different essential processes such as DNA replication, cell envelope synthesis, cytokinesis, as well as energy production to power these tasks. Additionally, carbon partitioning decisions should be made in conjunction with the assessment of nutrient availability and environmental threats such as those that weaken the cell envelope. GlmR (formerly YvcK) is a key metabolic factor involved in carbon utilization (**Fig. 1A**), specifically in the production of precursors needed for peptidoglycan (PG) synthesis and other cell envelope components (4–6). In this report, we examine the physiological impacts of GlmR, its absence, and the various regulatory means available to calibrate its function.

**Figure 1:**
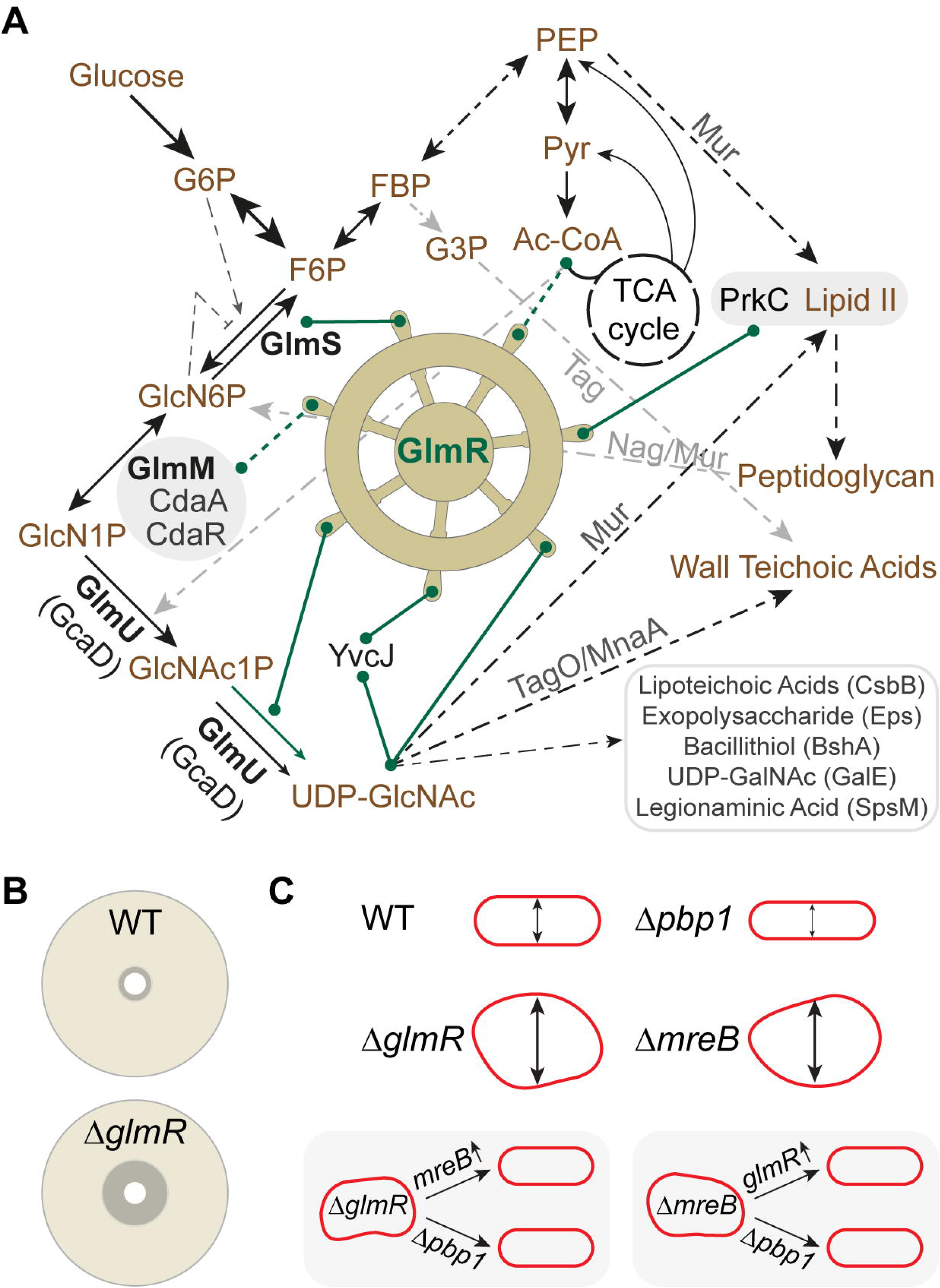
GlmR plays a central role in carbon metabolism, antibiotic resistance, and cell shape determination. **(A)** GlmR-centric metabolic pathway showing carbon flux in *B. subtilis*. Abbreviations: G6P, glucose-6-phosphate; F6P, fructose-6-phosphate; FBP, fructose 1,6-bisphosphate; G3P, glycerol-3-phosphate; PEP, phosphoenolpyruvate; Pyr, pyruvate; Ac-CoA, acetyl-coenzyme A; TCA, tricarboxylic acid; GlcN6P, glucosamine-6-phosphate; GlcN1P, glucosamine-1-phosphate; GlcNAc1P, N-acetylglucosamine-1-phosphate; UDP-GlcNAc, UDP-N-acetylglucosamine. The lines connected to GlmR wheel (green line) indicate direct (solid line) or indirect (dotted line) regulation and/or functions. Multistep pathways are shown in dotted black lines representing single or multiple steps respectively. Gray arrows indicate additional pathways not discussed in detail in this article. Processes/products that utilize UDP-GlcNAc are indicated with enzymes involved listed above the arrows or within parentheses. Enzymes needed to support the uridyltransferase activity (green arrow) of GlmR are shown in bold font. The catalytic activity of GlmR requires two consecutive aspartates at position 38 and 39. GlmR is acetylated at lysine 296 using Ac-CoA directly or indirectly with the help of another protein. GlmR is phosphorylated at threonine 304 by PrkC which is believed to sense Lipid II abundance. GlmR is regulated by UDP-GlcNAc binding which requires arginine 301. The function of GlmR in stimulating GlmS is modulated by another UDP-GlcNAc binding protein YvcJ. The genes *yvcJ* and *glmR* are encoded together. Evidence linking GlmR and the nucleotide second messenger cyclic-di-AMP (c-di-AMP) exists. *cdaA*-*cdaR*-*glmM* genes constitute a highly conserved operon encoding genes that form a tripartite complex which determines the intracellular concentration of c-di-AMP. *glmS* riboswitch-ribozyme activation/inhibition is indicated. **(B)** Diagrammatic summary of disk diffusion assay indicating cells lacking *glmR* are hypersensitive to cell wall targeting antibiotics. **(C)** Cartoon depiction of cell morphology of *B. subtilis* wild type (WT) and *glmR*, *pbp1* (*ponA*), and *mreB* deletion mutants. Deletion of *pbp1* results in thinner cells while deletion of *mreB* leads to increased cell width when viability is maintained via magnesium supplementation. Cells lacking *glmR* also are abnormally large when grown in gluconeogenic conditions. The cell morphology defects of both Δ*glmR* and Δ*mreB* can be corrected by either deletion of *pbp1* or by overexpression of *mreB* or *glmR* respectively.

*Antibiotic resistance:* As GlmR plays a key role in the accumulation of PG precursors, its presence becomes crucial in the presence of cell wall stress. Specifically, it has been noted that cells lacking *glmR* exhibit increased sensitivity to several cell wall targeting antibiotics (7, 8). This includes different classes of antibiotics that inhibit cell wall synthesis such as bacitracin, vancomycin, moenomycin, cefuroxime, and oxacillin.

Thus, GlmR is a critical antibiotic resistance factor (**Fig. 1B**).

*Cell shape:* It is known that GlmR becomes essential in conditions requiring gluconeogenesis (9). Cells lacking *glmR* grown in the absence of glucose exhibit abnormal morphology (**Fig. 1C**). Therefore, Δ*glmR* phenotypes can be suppressed by either glucose or magnesium supplementation (8, 9). While glucose would allow glycolysis and negate the need for gluconeogenesis, the effect of magnesium is likely multifactorial. This is because magnesium supplementation may inhibit PG hydrolases (10–12), deplete intracellular PG precursor levels (13), and facilitate osmoregulation (14). Besides chemical supplementation, deletion of *pbp1* (*ponA*), the gene encoding class A bifunctional penicillin binding protein (PBP1), also abrogates the cell morphology and viability defects of cells lacking *glmR* (**Fig. 1C**) (15). Additionally, overproduction of MreB, an actin-like cytoskeletal protein (16), also restores rod shape in Δ*glmR* strain (15). Conversely, overexpression of *glmR* or deletion of *pbp1* corrects the cell morphology defects of a strain harboring *mreB* deletion (15, 17). Balanced activities of MreB and PBP1 have been recognized to govern the width of the cell (18, 19). Given the reciprocal phenotypes of *mreB* and *glmR* mutants, it is possible GlmR is also a cell width determining factor. In this report we present evidence in support of this notion.

*Cell wall precursor synthesis:* The first step to commit carbon for the PG precursor pathway is taken by GlmS, an enzyme involved in the de novo synthesis of GlcN6P (key to abbreviations are provided in **Fig. 1** legend). *glmS* transcript abundance is self-regulated through autocleavage by a GlcN6P-responsive riboswitch-ribozyme (20). This ribozyme activity is hindered by glucose and G6P (21, 22). Intriguingly, changes in intracellular pH and magnesium concentration may influence the activity of this ribozyme (23, 24). Alternatively, peptidoglycan can be recycled into GlcN6P to supplement this pathway (25). Subsequently the enzymatic actions of GlmM and GlmU (GcaD) help generate the essential cell wall precursor, UDP-GlcNAc. In *B. subtilis*, GlmR directly interacts with GlmS and stimulates its enzymatic activity (8, 26, 27).

Moreover, it was shown recently that GlmR of *B. subtilis* and other species are in fact uridyltransferase enzymes capable of producing UDP-GlcNAc (27). Therefore, GlmR plays a central role in carbon utilization by influencing the accumulation of UDP-GlcNAc (**Fig. 1A**). Consequently, the flow of carbon to make PG precursors can be calibrated by regulating GlmR.

*Nucleotide second messenger signaling:* Past studies have identified mutations that bypass the need for *glmR* to increased expression of *glmM* and/or *glmS* (8). This genetic locus harbors the highly conserved *cdaA*-*cdaR*-*glmM* operon (**Fig. 1A**) (28). CdaA is the major cyclic-di-AMP (c-di-AMP) synthase and is regulated by CdaR and GlmM in *B. subtilis* and other organisms (29–33). Intriguingly, disruption of c-di-AMP phosphodiesterase genes *pgpH* (*yqfF*) or *gdpP* (*yybT*) also alleviate *glmR* phenotypes (8, 9). Yet, the mechanism behind this remains unclear.

*Post-translational regulation:* GlmR binds UDP-GlcNAc, the product of its catalytic activity (**Fig. 1A**). When UDP-GlcNAc level is in excess, this ligand binding is thought to weaken the GlmR-mediated stimulation of GlmS to establish a negative feedback loop (26, 34). Additionally, YvcJ, a RapZ-like protein encoded from the gene immediately upstream of *glmR*, also binds UDP-GlcNAc and GlmR to moderate the function of the latter (26). Besides ligand binding, GlmR (T304 residue) is also subject to phosphoregulation by PrkC (7), a S/T kinase speculated to regulate cell wall homeostasis by sensing the level of Lipid II precursors (35–37). Furthermore, a proteomics study identified that GlmR (K296 amino acid) is acetylated (38). Metabolic intermediates such as acetyl-CoA and its derivative acetyl-phosphate are known to serve as donors of acetyl group for lysine acetylation (39, 40).

In this study, we sought to investigate the new and unresolved questions regarding GlmR in antibiotic resistance, cell morphogenesis, and c-di-AMP signaling. In addition, we probed the significance of enzymatic activity, UDP-GlcNAc binding, phosphorylation, and acetylation of GlmR. Our results reveal that GlmR aids in resisting tunicamycin, an antibiotic that targets wall teichoic acids (WTA) biosynthesis selectively and the PG pathway at higher concentrations. We observe that cells lacking GlmR exhibit reproducibly larger cell width compared to WT control even when irregular cell morphology is corrected with chemical supplementation. Reciprocally, we see a reduction in cell width with *glmR* overexpression. We notice that division site positioning is impaired when *glmR* is deleted. We also note that c-di-AMP level is elevated in the absence of GlmR in a manner dependent on CdaA. Our experiments demonstrate that the uridyltransferase activity of GlmR is essential for its cellular function, as the catalytically inactive mutant is unable to complement the Δ*glmR* phenotypes. Finally, we show that UDP-GlcNAc binding, phosphorylation, and lysine acetylation finetune GlmR activity. We provide a model based on our results to explain the role of GlmR in reversing the phenotypes of cells lacking actin-like proteins MreB and Mbl. In sum, our results shed light on the multi-level regulation of GlmR and its crucial role in cell morphogenesis and antibiotic resistance.

## RESULTS

### GlmR is important for tunicamycin resistance

Given the essentiality of UDP-GlcNAc in WTA production (**Fig. 1A**) (41), we reasoned that cells lacking GlmR may display increased susceptibility to antibiotics that target the WTA pathway, such as tunicamycin (**Fig. 2**). The nucleoside antibiotic, tunicamycin, is an analog of UDP-GlcNAc, is known to inhibit glycosyltransferases (42). More specifically, the first step of WTA biosynthesis mediated by TagO is selectively inhibited by tunicamycin at a lower concentration range, while at a significantly higher range (>100x concentration in *S. aureus* (43); >50 µg/ml in *B. subtilis* (44, 45)) hinders the function of MraY involved in the PG synthesis pathway (**Fig. 2C**) (45–47). To test whether there are any changes in tunicamycin susceptibility between WT and cells lacking *glmR*, we conducted a disk diffusion assay encompassing a range of concentrations. Briefly, sterile disks laced with 0, 10, 25, 50, and 100 µg/ml of tunicamycin were placed on the lawns of either WT or Δ*glmR* strains (**Fig. 2A**). The WT strain did not exhibit any noticeable growth defect at 10 µg/ml and we noticed a smaller zone of inhibition (ZOI) at 25 µg/ml. Remarkably, consistent with our prediction, we observed a larger ZOI for Δ*glmR* compared to WT control at all concentration ranges including 10 µg/ml, the lowest concentration tested (**Fig. 2AB**; see red arrows). The increased susceptibility of Δ*glmR* was abolished with *glmR* complementation, even in the absence of inducer likely due to leaky expression. As GlmR plays a key role in supplying UDP-GlcNAc (**Fig. 1A**) for both TagO and MraY enzymes, it is conceivable that the biogenesis of both WTA and PG (the two most important cell envelope components (48)) are dysregulated in the absence of GlmR. Thus, cells lacking *glmR* are ill-equipped to counter antibiotics that target these pathways. Based on our results, we add tunicamycin to yet another class of antibiotics for which resistance is conferred by the presence of GlmR.

**Figure 2:**
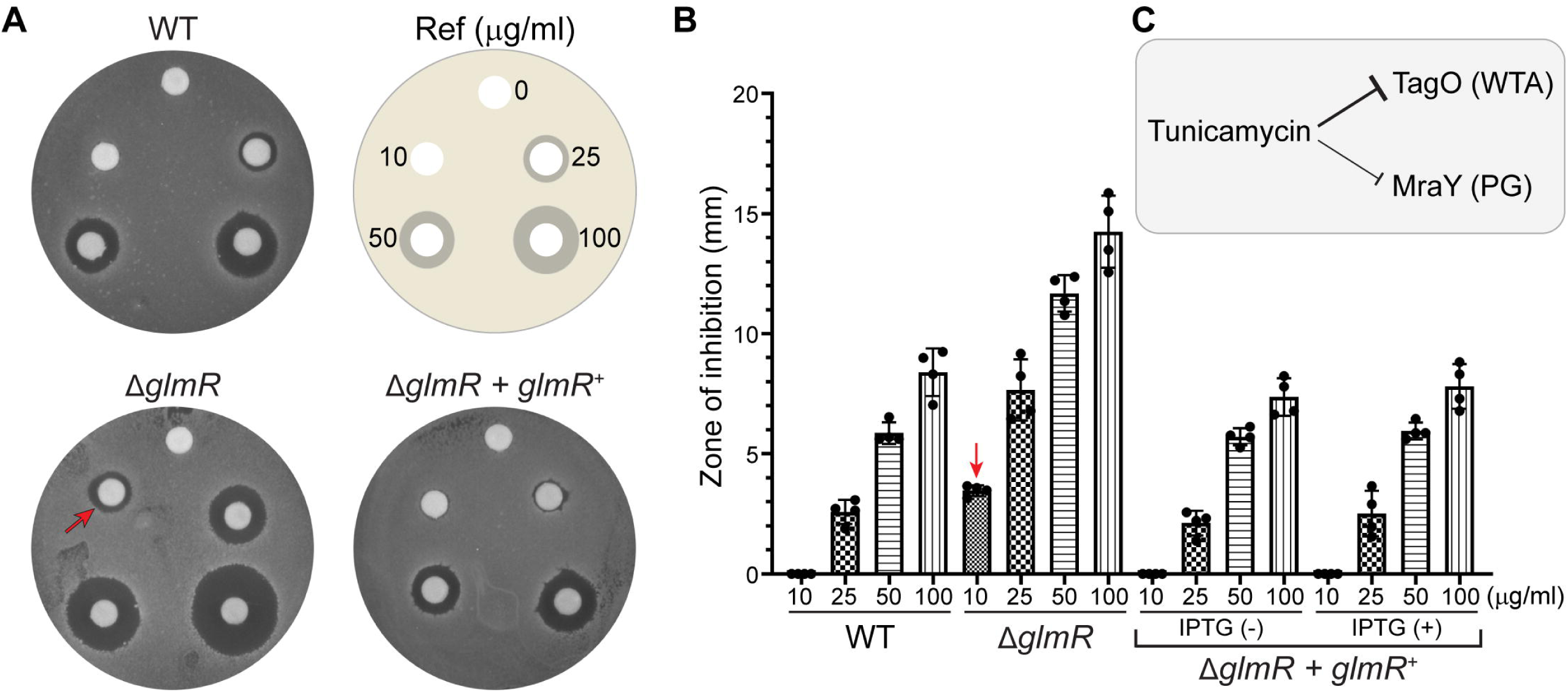
Cells lacking *glmR* exhibit increased tunicamycin sensitivity. **(A)** Discs containing 5 µl of either 0, 10, 25, 50, or 100 µg/ml tunicamycin were placed (as shown in reference; Ref) on lawns made of WT (PY79), Δ*glmR* (SK35), and Δ*glmR* complemented with IPTG-inducible *glmR* (SK56). Absence of GlmR leads to increased tunicamycin susceptibility, as evidenced by increased zone of inhibition (ZOI) compared to WT and the complementation strain. **(B)** Quantification of the ZOI (minus the 6.5 mm disc) of strains shown in panel A. Also plotted are data for SK56 strain grown in the absence (leaky expression) or presence (1 mM) of IPTG. Average of four replicates is shown, and standard deviation is displayed as error bars. The red arrow indicates the concentration where ZOI was observed for Δ*glmR* but not for WT. **(C)** Enzymes targeted by tunicamycin. TagO, an enzyme critical for synthesizing wall teichoic acids (WTA) is selectively inhibited at lower concentration range while MraY involved in peptidoglycan (PG) synthesis is also inhibited at higher concentrations.

### GlmR is required for cell width maintenance

Given that *glmR* and *mreB* mutants phenocopy each other (**Fig. 1C**), we tested whether GlmR is also involved in cell width maintenance similar to MreB. As mentioned earlier, it is known that glucose or magnesium supplementation supports Δ*glmR* cell viability in conditions where GlmR is otherwise essential (9). Thus, we examined the morphology of wild type (WT) and Δ*glmR* cells using fluorescence microscopy with or without glucose and magnesium supplementation in lysogeny broth (LB). We quantified the cell morphology changes for 4 h to cover both exponential growth phase and transition to stationary phase (**Fig. 3A**). As expected, Δ*glmR* cells exhibited abnormal cell bulging compared to WT control in the absence of any supplementation (**Fig. S1A**). We also observed increased incidence of aberrant division site positioning in the Δ*glmR* strain with nearly 40% of septa in the field of view (n=100) when compared to WT (<5%; n=100) (**Fig. 3B** and **Supplementary Movie 1**). Thus, it appears that proper positioning of cytokinetic machinery is impaired when GlmR is absent.

**Figure 3:**
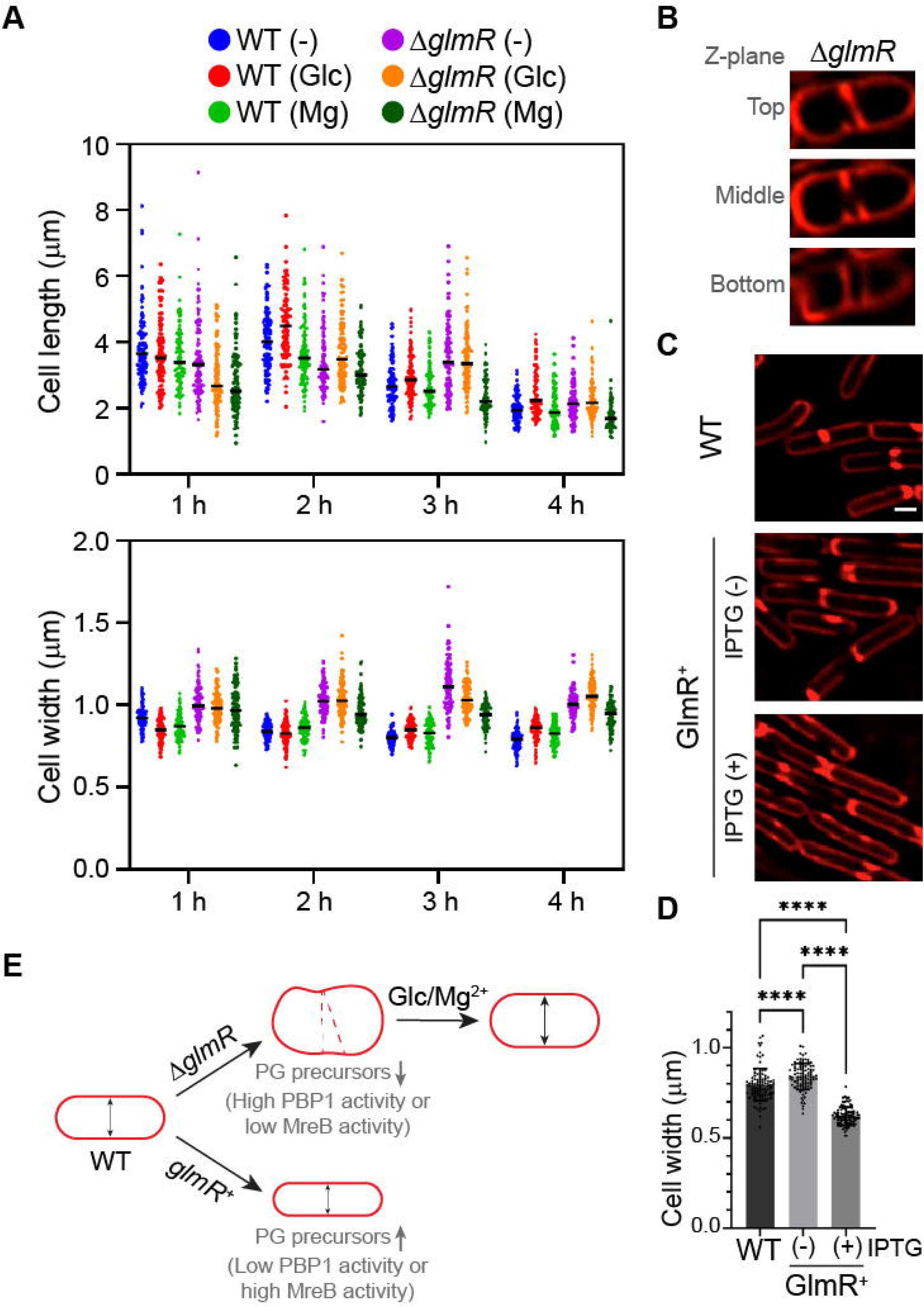
GlmR is a cell width determinant. **(A)** Cell length and width quantification of WT (PY79) and Δ*glmR* (RB176) strains grown in LB in the absence or presence of D-glucose (1%) or magnesium (25 mM MgCl_2_) supplementation grown for up to 4 hours and imaged at every hour. Representative micrographs are shown in Fig. S1A. Individual data points and corresponding average (black line) of indicated time points post-supplementation or mock are shown (n = 100). **(B)** Micrographs showing abnormal septation in Δ*glmR* (SK35). See **Supplementary Video 1** to view all frames. Additional examples are indicated in Fig. S1 with yellow arrows. Red, FM 4-64 membrane dye. **(C)** Representative micrographs showing WT (PY79) and Δ*glmR glmR^+^*(SK56) uninduced or induced with 1 mM IPTG. Scale bar, 1 µm. **(D)** Cell width quantification of panel C data (n = 100). Cells were measured using FIJI. Statistical significance was assessed through one-way ANOVA with Tukey’s correction; **** = p<0.0001. **(E)** Graphical summary depicting key observations. Although glucose or magnesium supplementation abrogates the abnormal morphology of Δ*glmR* cells, the average cell width is consistently larger than WT control at all time points. Additionally, we note that overexpression of *glmR* leads to reduced cell width and that deletion of *glmR* impairs division site positioning indicated as red dotted lines.

The cell length changes over time were consistent with previous observations for WT (**Fig. 3A**) (49, 50). A similar trend was seen for the cells lacking GlmR. Δ*glmR* cells at 3 h timepoint were longer than the WT control, which was noted previously (34). However, by 4 h mark this difference was diminished. In contrast to cell length, the cell width difference between WT and Δ*glmR* was more striking – with the latter being consistently wider. Magnesium addition corrects the abnormal Δ*glmR* cell morphology as reported previously (9). We find that magnesium supplementation leads to shorter cell length at nearly all time points regardless of the strain type. This specific effect of magnesium on *B. subtilis* cell length has been reported previously (51). Nonetheless, the cell width is consistently larger than WT. Although the aberrant cell morphology in the absence of GlmR is known, we expected glucose supplementation to allow glycolysis (as GlmR function is believed to be prominent only during gluconeogenesis (15)) and restore cell width similar to that of WT. However, our results reveal that Δ*glmR* cells are wider than WT at all timepoints even in the presence of glucose. Likewise, the post-exponential phase cell widths at 4 h were also greater than the WT control with or without supplementation. Reciprocally, unlike deletion, overexpression of *glmR* which presumably increases the abundance of PG precursors, leads to decreased cell width (**Fig. 3CD**). Based on these results, we infer that GlmR is a key cell width determinant (**Fig. 3E**), possibly through assisting PBP1 localization (15) and/or by feeding enough PG precursors to balance MreB/PBP1 consumption (**Fig. 1C**).

### Absence of GlmR leads to CdaA-dependent elevation in c-di-AMP level

In vegetatively growing *B. subtilis*, c-di-AMP synthesis occurs mainly through CdaA and DisA, which are the major and minor synthases respectively (**Fig. 4A**) (8, 32, 52, 53). It is known that cells lacking both are nonviable (54, 55), thus highlighting the vital role of c-di-AMP signaling. Given that mutations in the *cdaA* locus or deletion of c-di-AMP hydrolases allow Δ*glmR* phenotype suppression (8, 9, 56), we wished to probe the intracellular c-di-AMP level in cells lacking *glmR*. For this, we engineered a reporter in which *gfp* expression was controlled by the c-di-AMP sensing riboswitch of *kimA* (*ydaO*) promoter (57–59). Briefly, when the intracellular level of c-di-AMP drops, *gfp* will be transcribed and when c-di-AMP level is high *gfp* expression will halt (**Fig. 4B**).

**Figure 4:**
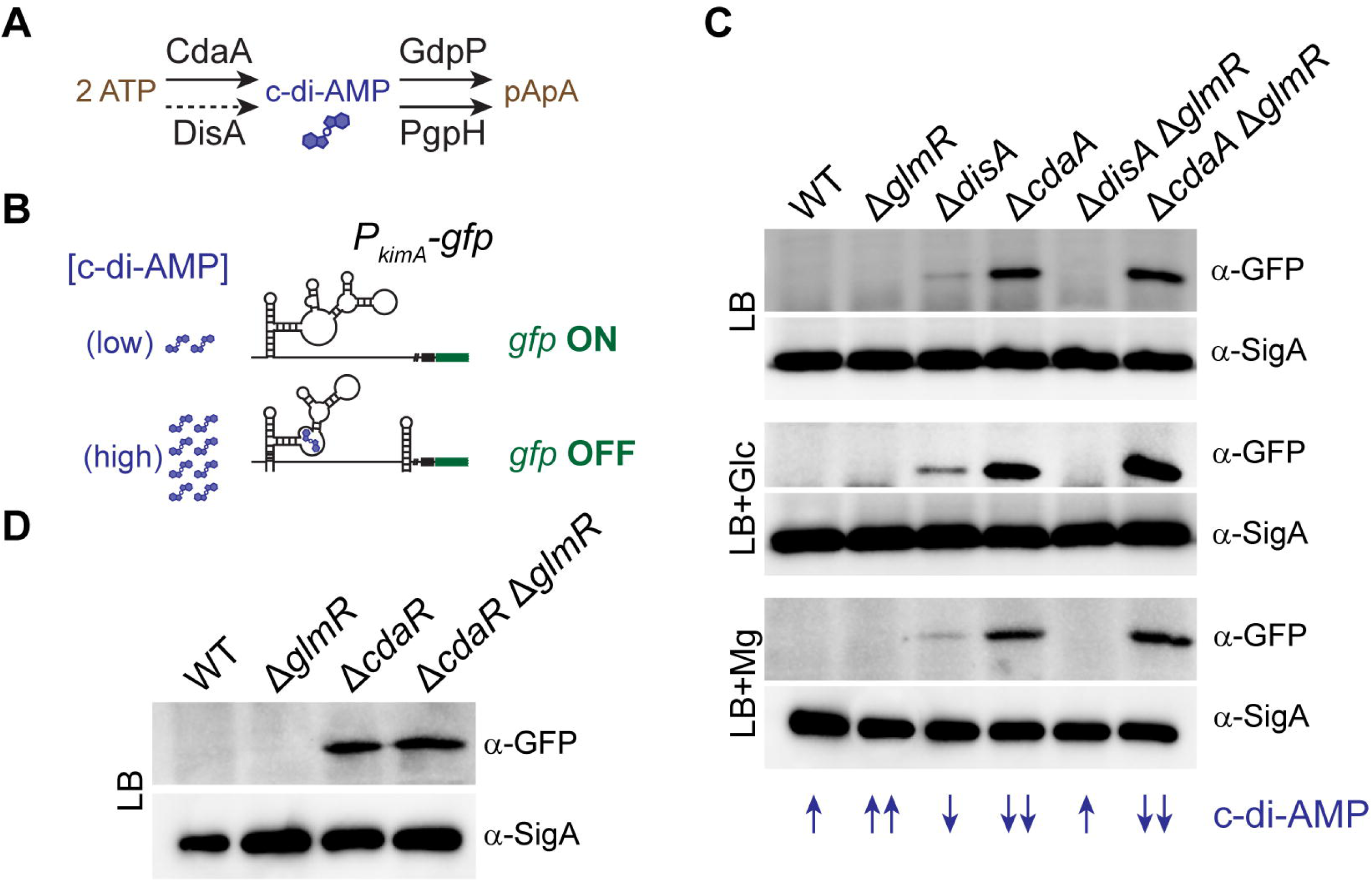
Absence of GlmR stimulates c-di-AMP production. **(A)** Enzymes involved in the c-di-AMP synthesis and turnover. CdaA and DisA are the major and minor c-di-AMP synthases respectively. *B. subtilis* encodes a third c-di-AMP synthase, CdaS, that is activated specifically during sporulation (not shown here). GdpP and PgpH are phosphodiesterases involved in the conversion of c-di-AMP to phosphoadenylyl-adenosine (pApA). Solid lines and dotted lines indicate the major and minor enzymes respectively. **(B)** C-di-AMP riboswitch reporter of *kimA* was used to generate a GFP-based reporter. Transcription of *gfp* is dependent on low intracellular c-di-AMP concentration. **(C)** Representative western blots of *P_kimA_-gfp* reporter in WT (SK94), Δ*glmR* (SK113), Δ*disA* (SK109), Δ*cdaA* (SK110), Δ*disA* Δ*glmR* (SK111), and Δ*cdaA* Δ*glmR* (SK112) backgrounds. All strains were grown in LB without or with supplementation of D-glucose (1%) or MgCl_2_ (25 mM) and harvested 2 hours after induction. Antibodies against GFP and SigA (loading control) were used to monitor the changes in *P_kimA_-gfp* reporter activity. Blue arrows indicate inferred changes in c-di-AMP levels as discussed in the results section relative to WT (↑). **(D)** Immunoblot of WT (SK94), Δ*glmR* (SK113), Δ*cdaR* (SK118), and Δ*glmR* Δ*cdaR* (SK119) probed with GFP or SigA antisera.

Therefore, based on our design, lack of GFP indicates presence of c-di-AMP and vice versa. Similarly designed reporters have been successfully utilized for this purpose by other groups (60, 61). We introduced this reporter in the WT background as well as in strains harboring individual or multiple deletions of *glmR*, *cdaA*, and *cdaR*. Previous reports have found that relative to WT, deletion of *disA* leads to a modest decrease in intracellular c-di-AMP concentration (61–64). As our probe lacks the ability to distinguish high vs. very high levels of c-di-AMP, to enhance the sensitivity of our reporter, we also included *disA* deletion. In WT and Δ*glmR*, we did not detect a GFP band suggesting these cells maintain relatively high intracellular c-di-AMP concentration (**Fig. 4C**). As expected, cells lacking *disA* produced a faint band while the *cdaA* knockout strain produced an intense band. Interestingly, in the strain harboring deletions of both *disA* and *glmR* genes, the faint GFP band was no longer present (**Fig. 4C**; compare Δ*disA* and Δ*disA* Δ*glmR* lanes). This trend was consistent and reproducible even when the respective cultures were grown in the presence of glucose or magnesium. We infer this to mean that lack of GlmR leads to stimulation of c-di-AMP production. We do not observe any noticeable changes in *cdaA glmR* double deletion as the GFP band intensity corresponding to this strain resembles that of *cdaA* knockout. Thus, the increased c-di-AMP levels we notice in Δ*disA* Δ*glmR* strain appears to be dependent on CdaA. Next, we tested whether this stimulation requires CdaR, a known modulator of CdaA activity (32, 55). Deletion of both *cdaR* and *glmR* also produced a prominent GFP band similar to Δ*cdaA* and Δ*glmR* (**Fig. 4D**). Our results suggest that both CdaA and CdaR are responsible for the elevated c-di-AMP levels in the Δ*disA* Δ*glmR* strain.

Therefore, absence of GlmR leads to stimulation of c-di-AMP production through the major c-di-AMP synthase CdaA.

Inspired by this observation, we probed whether the changes in c-di-AMP levels influence the Δ*glmR* cell morphology. We find that deletion of both *cdaA* and *glmR* results in a stunning reversal of aberrant cell morphology (**Fig. S2A**). However, as discussed in the supplemental information, we attribute this to a polar effect that may synthetically elevate the expression of the downstream gene *glmM* of the *cdaA*-*cdaR*-*glmM* operon (**Fig. S2BC**) (65, 66). Notably, suppression of Δ*glmR* phenotypes by increased levels of GlmM has been reported previously (8, 56). Regardless, we note that deletion of *cdaA* or *cdaR* individually or in combination with Δ*glmR* results in a significant drop in intracellular c-di-AMP level (**Fig. 4CD**) - yet they all maintain normal cell shape (**Fig. S2A**). Based on this, we conclude that reduced intracellular c-di-AMP concentration is not quite detrimental to cells when the cell envelope is not compromised or stressed.

We also tested whether other c-di-AMP signaling related enzymes indicated in **Fig. 4A** influence the Δ*glmR* phenotypes. For this, we investigated the double deletion phenotypes of a strain devoid of *glmR* and the gene encoding the minor c-di-AMP synthase DisA. We also included the strains lacking GlmR and one of the two c-di-AMP phosphodiesterases GdpP or PgpH. Our experiments show that additional deletion of *disA*, *gdpP*, or *pgpH* is unable to correct the abnormal Δ*glmR* cell shape defect (**Fig. S2AD**). However, it appears to partially correct the growth phenotypes to a varying degree (**Fig. S3**). Our interpretations of these results and associated phenotypes are discussed in the supplemental file.

### The enzymatic activity of GlmR is integral for its function

GlmR was recently shown to be an uridyltransferase (27). Therefore, we aimed to probe the physiological relevance of the enzymatic function of GlmR. To examine this, we generated strains in which Δ*glmR* is complemented with an inducible copy of either unmutated *glmR* or *glmR* variant harboring catalytic site mutations (D38A D39A; (27)). We then tested the growth of these strains on LB agar (LA) and Difco starch (DS) plates (**Fig. 5A**). On LA (which contains magnesium (51)), when compared to WT, Δ*glmR* cells grew, albeit poorly. In contrast, on DS, while the WT strain grew well, cells lacking *glmR* exhibited severe growth inhibition and appear to be nearly non-viable. This medium is similar to Mueller-Hinton (MH) which is prohibitive for supporting the growth of Δ*glmR* strain (8). However, we are unsure why Δ*glmR* cells are unable to utilize starch to support glycolysis. It is possible defective WTA may hinder the secretion of amylase needed for starch degradation (67). Nevertheless, both the small colony phenotype on LA and lethality on DS can be reversed by *glmR* complementation. Thus, DS growth medium serves as a useful tool to study the functionality of GlmR mutants.

**Figure 5:**
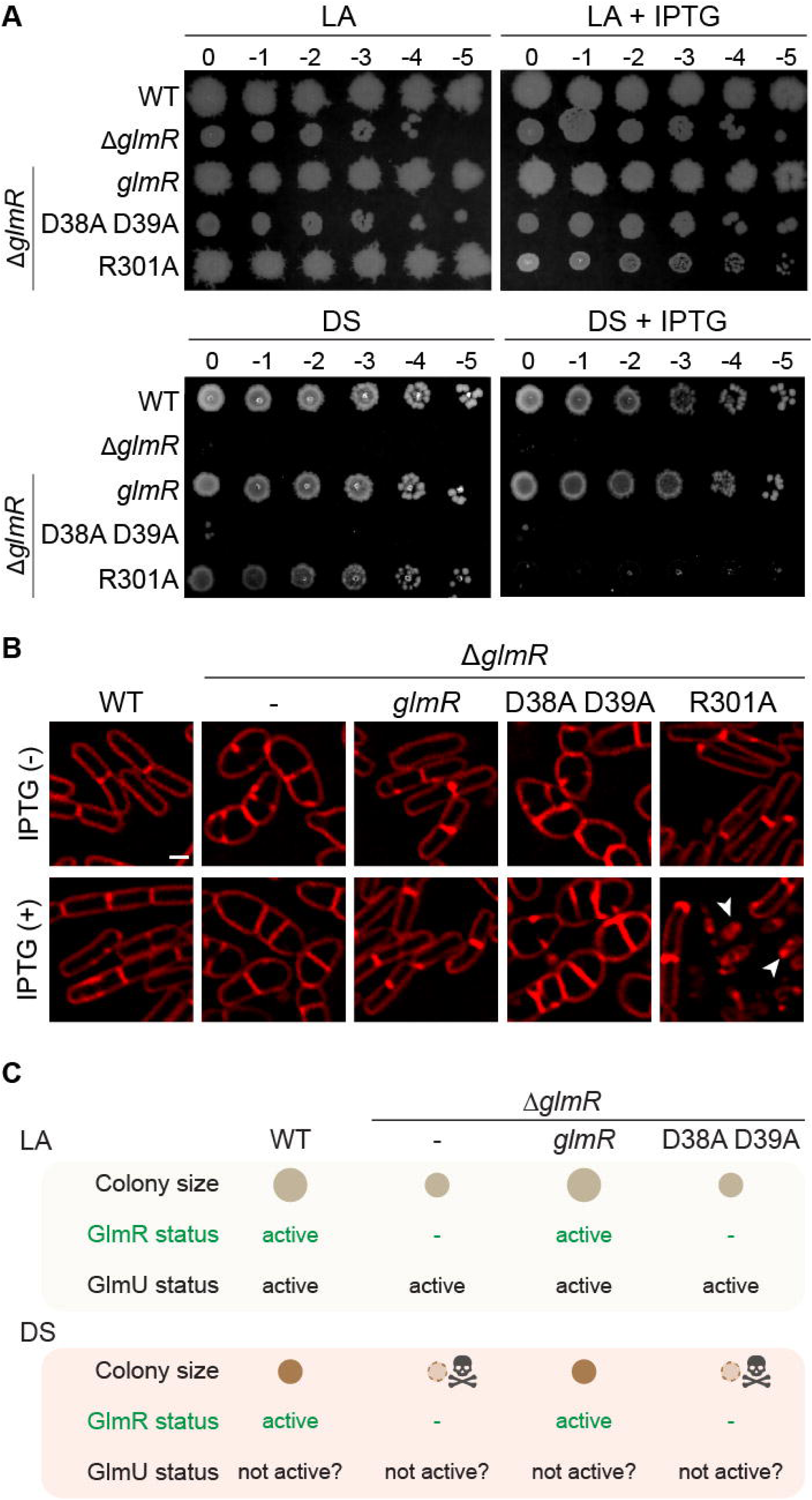
Enzymatic activity of GlmR is essential for its function. **(A)** Serial dilutions of WT (PY79), Δ*glmR* (RB176) and Δ*glmR* complemented with IPTG-inducible *glmR* (SK56), D38A D39A *glmR* mutant that is catalytically inactive (BLS95), or R301A mutant which is deficient in UDP-GlcNAc binding (SK141). Representative pictures of LA and DS plates incubated overnight at 37 °C are shown. Inducer plates contain IPTG (1 mM) when indicated. **(B)** Micrographs of strains listed in panel A grown in LB without or with (1 mM) IPTG. Cells are labeled with FM 4-64 membrane dye (red). Scale bar, 1 μm. Arrow heads point to lysed cells. **(C)** Tabulated summary of data shown in panel A indicating the functionality of GlmR and speculated activity of GlmU in the indicated strain backgrounds and growth media.

Interestingly, we find that the D38A D39A mutant is unable to complement the Δ*glmR* phenotypes on LA and DS (**Fig. 5A**). Cell morphology analysis through microscopy reveals that D38A D39A mutant is not capable of correcting the cell shape abnormality commonly seen in Δ*glmR* strain (**Fig. 5B**). We used functional His-tagged GlmR to ensure that the D38A D39A mutant is stably produced (**Fig. S4**). Our results reveal that the catalytic activity of GlmR is required for complementation and therefore for its normal cellular function. GlmU is the primary (essential) enzyme responsible for UDP-GlcNAc production (**Fig. 1A**). As depicted in **Fig. 5C**, combined action of GlmU and GlmR is likely needed to support WT-like growth on LA. Thus, we see poor growth without the enzymatic function of GlmR. On the other hand, on DS medium, our results indicate that GlmR becomes the sole enzyme responsible for UDP-GlcNAc production. This notion is further strengthened by our finding that the catalytically inert mutant is unable to support the growth of Δ*glmR* strain.

Next, we tested whether UDP-GlcNAc binding by GlmR affects growth by making use of the R301A mutant that lacks the ability to bind to this ligand. As previously reported (8, 34), R301A complements Δ*glmR* phenotypes. However, we note this happens on LA and DS only in the absence of inducer due to leaky expression - unlike the catalytically inert mutant (**Fig. 5A**). However, in the presence of inducer, the phenotype is starkly different. We noticed a decrease in colony diameter on LA + IPTG and no growth on DS + IPTG. Thus, overexpression of R301A closely mimics Δ*glmR* phenotype in these conditions. Upon probing these cells through microscopy, we find that the cells harboring an inducible copy of R301A mutant are more similar to WT in the absence of inducer and prone to lysis in the presence of inducer (**Fig. 5B**; see arrowheads). UDP-GlcNAc binding is proposed to promote GlmR-YvcJ interaction and thereby prevent GlmR-mediated stimulation of GlmS (**Fig. 1A**) (26). Therefore, it is possible that the R301A mutant may continuously stimulate GlmS which is detrimental to cell viability. Although the differential effect on YvcJ phenotype for this mutant has been noted (26), the toxic effect upon overexpression was not. This data highlights the regulatory potential of UDP-GlcNAc ligand – which is the product of the enzymatic activity of GlmR.

### GlmR phosphorylation may negatively affect UDP-GlcNAc binding

GlmR is a substrate of the S/T kinase PrkC and is phosphorylated at T304 (7). Therefore, we tested the ability of T304A (phosphoablative) and T304E (phosphomimetic) versions of GlmR to complement Δ*glmR* phenotypes. Our results indicate that both mutants function equivalently to unmutated *glmR* and complement all *glmR* knockout phenotypes: growth on LA and DS (**Fig. S5A**) and restoration of cell morphology (**Fig. S5B**). We also note that similar to *glmR* overexpression (**Fig. 3CD**), overproduction of phosphomimetic or phosphoablative versions of GlmR results in cell width reduction (**Fig. S5C**). Thus, phosphorylation does not appear to significantly influence GlmR function. This observation is consistent with previous reports (7, 8).

Upon closer inspection, we do notice a modest but reproducibly smaller colony size for T304A phosphoablative mutant when compared to the GlmR control and T304E variant (**Fig. S5D**). It has been noted that GlmR phosphomutants have differential effects on bacitracin sensitivity, correction of aberrant cell shape of Δ*mreB*, and PBP1 localization (7). Thus, we aimed to further investigate the specific purpose of GlmR phosphoregulation.

Given the proximity of T304 phosphosite to R301, the residue important for UDP-GlcNAc binding, GlmR phosphorylation may influence ligand recognition (34).

Therefore, we tested this possibility with purified GlmR. For this, we used FITC-labeled protein (WT GlmR or mutants) and monitored the fluorescence change associated with ligand (UDP-GlcNAc) binding to study the binding kinetics (68). The saturation of change in fluorescence (ΔF) associated with UDP-GlcNAc titration allowed us to estimate the dissociation constant (K_d_) for GlmR as 0.28 ± 0.04 mM (**Fig. 6A**; see inset). This value is close to 0.41 mM predicted previously (34), highlighting the reliability of this approach. In contrast, for our negative control R301A mutant, we did not notice any change in fluorescence. This confirms that R301A mutant is unable to bind UDP-GlcNAc as established previously (34). For T304A mutant, our estimated K_d_ value is 1.67 ± 0.31 mM, which is in a similar range to GlmR control. On the contrary, for T304E mutant, ΔF saturation was not achieved even with 5 mM UDP-GlcNAc, which precluded us from calculating the K_d_ value. We infer this result to mean that UDP-GlcNAc binding is possibly weaker and/or more transient when GlmR is in a phosphomimetic state (**Fig. 6B**). Overall, our results indicate that GlmR phosphorylation may render UDP-GlcNAc binding less favorable (**Fig. 6C**). As PrkC is involved in the regulation of cell envelope synthesis and is important for resisting cell wall targeting antibiotics (35–37), GlmR phosphorylation may be used by cells to override self-inhibition by UDP-GlcNAc accumulation to accelerate PG precursor production and strengthen the cell envelope.

**Figure 6:**
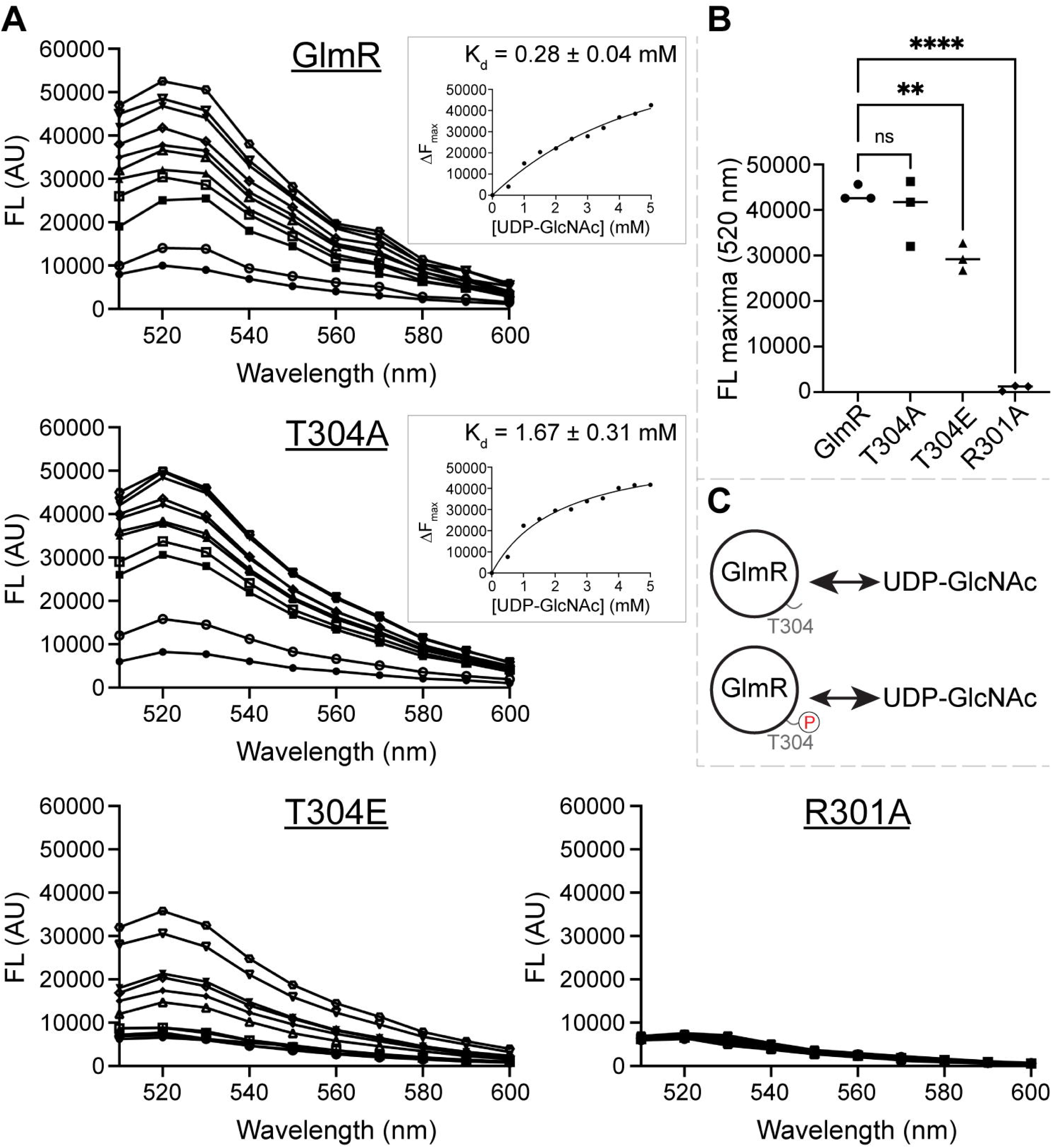
Phosphorylation of GlmR may weaken UDP-GlcNAC binding. **(A)** The binding of UDP-GlcNAc to purified WT GlmR (SK56), GlmR T304A (SK139), GlmR T304E (SK140), or GlmR R301A (SK141) were monitored using fluorescence spectroscopy. Shown are the representative fluorescence spectra of FITC-labeled GlmR or mutants (5 µM) were incubated without (●) or with UDP-GlcNAc at 0.5 mM (○), 1.0 mM (▪), 1.5 mM (□), 2.0 mM (▲), 2.5 mM (△), 3.0 mM (⍰), 3.5 mM (⍰), 4.0 mM (▾), 4.5 mM (⍰) and 5.0 mM (⍰) concentrations. Inset, binding constants (K_d_) were calculated by plotting the difference in fluorescence intensities (ΔF_max_) at different ligand concentrations when saturation was achieved. **(B)** The fluorescence maxima of GlmR (●), T304A (▪), T304E (▲), and R301A (⍰) are plotted; n=3, ** and **** indicate p <0.0084 and 0.0001, respectively. **(C)** Inference model based on the results shown in panel A indicating that phosphoregulation of residue T304 may alter the kinetics of UDP-GlcNAc ligand binding.

### GlmR function is likely affected by acetylation

In addition to phosphorylation, GlmR (also known as MgfK) is subject to another post-translational modification through lysine acetylation at position 296 (38). Several other metabolic enzymes shown in **Fig. 1A**, including GlmS, GlmM, GlmU, and YvcJ are also acetylated (38, 39, 69, 70). Acetylation is known to regulate catalysis, protein structure, and alter partner preference between proteins (71). Therefore, we aimed to study whether lysine acetylation influences GlmR function. To investigate this, we mutated K296 to either glutamine to mimic acetylation or arginine to resemble an unacetylated residue as previously described (38). As shown in **Fig. 7A**, K296Q acetyl-mimetic mutant was able to complement Δ*glmR* phenotypes in LA and DS similar to the *glmR* control. In contrast, while K296R acetyl-ablative mutant was able to complement in the absence of inducer, overexpression resulted in significant growth inhibition (**Fig. 7A**; see + IPTG plates). Microscopy analysis revealed that K296Q acetyl-mimetic mutant corrected the abnormal Δ*glmR* cell morphology, similar to the unmutated *glmR* control (**Fig. 7B**). On the other hand, K296R acetyl-ablative mutant prevented the cell bulging phenotype of Δ*glmR* only in the absence of inducer. In the presence of inducer, we observed increased cell lysis (**Fig. 7B**; see arrowheads). Thus, it appears that acetylation may also moderate the activity of GlmR by possibly hindering UDP-GlcNAc binding as these observations resemble the phenotypes of R301A variant (**Fig. 5**).

**Figure 7:**
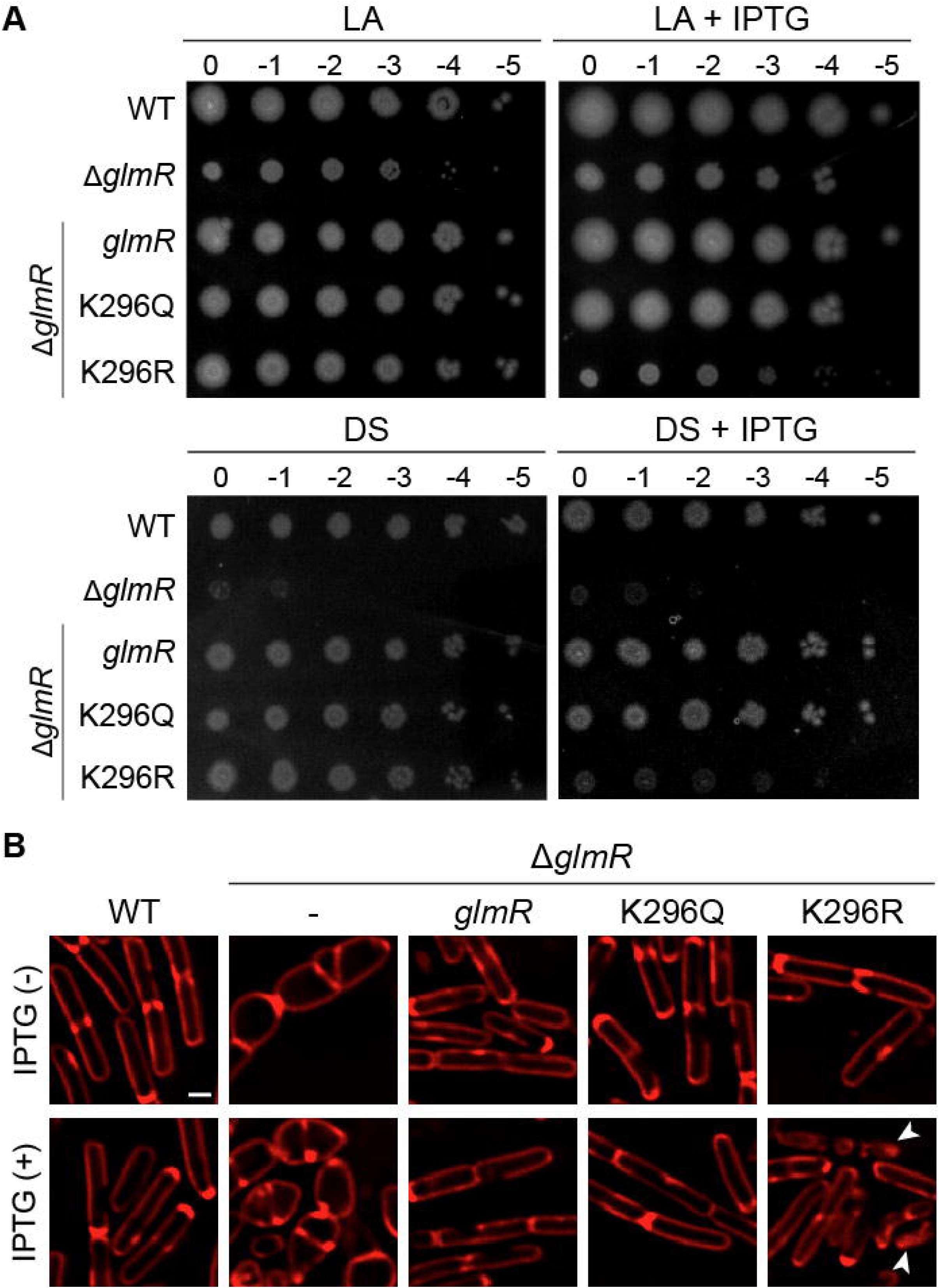
GlmR function is moderated by acetylation. **(A)** Spot titers of WT (PY79), Δ*glmR* (RB176) and Δ*glmR* complemented with IPTG-inducible *glmR* (SK29), K296Q *glmR* mutant to mimic acetylation (SK107), or K296R *glmR* mutant to mimic unacetylated status (SK108). Representative plate pictures of the above-mentioned strains grown in LA or DS medium at 37 °C overnight are shown. Inducer plates contain IPTG (1 mM). **(B)** Fluorescence micrographs of strains listed in panel A grown in LB in the absence or presence IPTG (1 mM). Cell membrane is stained with FM 4-64 dye (red). Scale bar, 1 μm. Arrowheads indicate cell lysis.

Taking the proximity of the sites important for UDP-GlcNAc binding (R301) and phosphorylation (T304) and our results into account, we propose that acetylation of GlmR (K296) likely negatively affects the flow of carbon to the PG precursor pathway. Conversely, we believe deacetylation and phosphorylation may promote UDP-GlcNAc synthesis (**Fig. 8A**).

**Figure 8:**
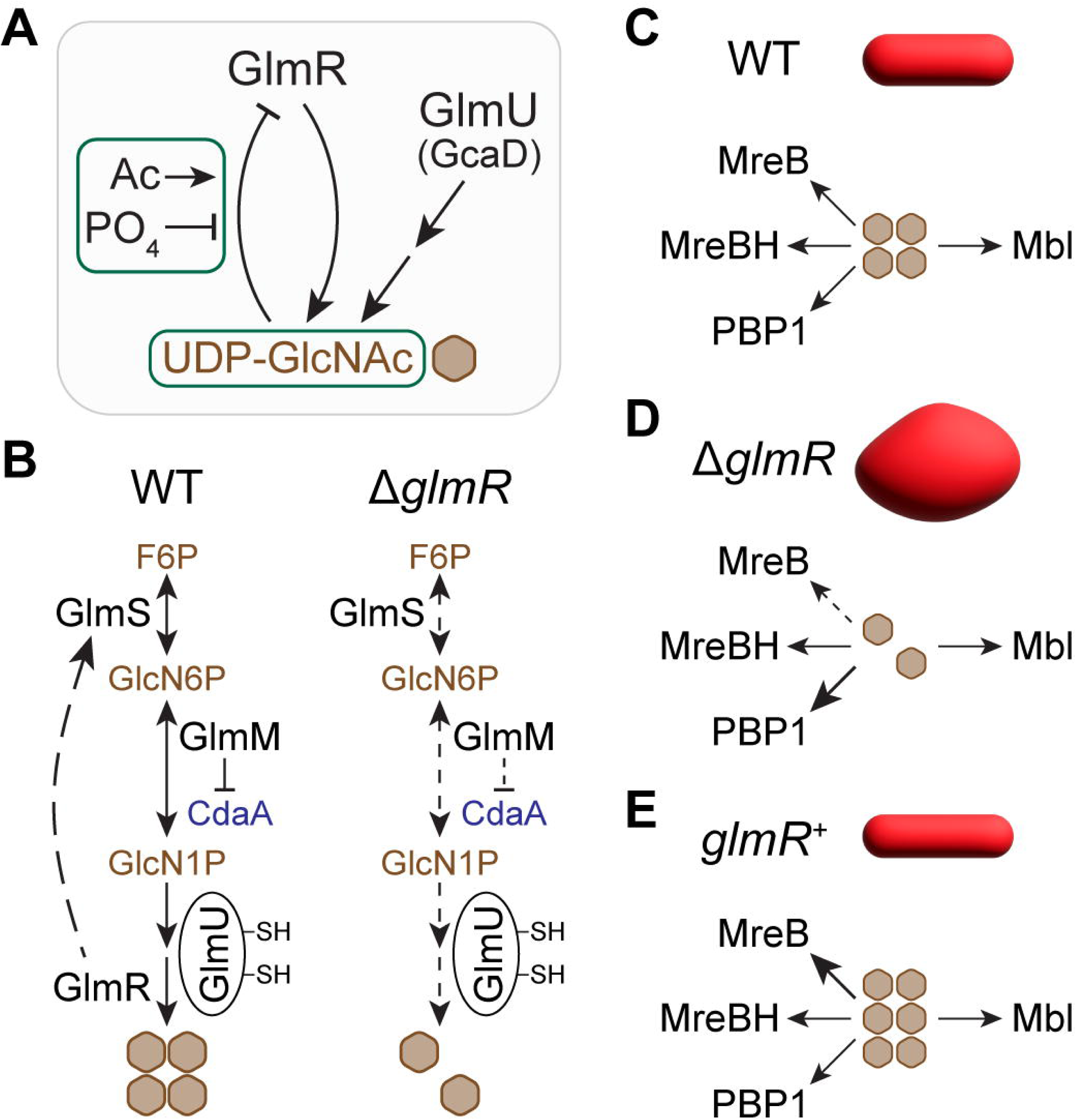
Graphical summary and illustration of proposed models. **(A)** In *B. subtilis,* GlmU is the main bifunctional essential enzyme that generates UDP-GlcNAc. In this report, we show that the uridyltransferase activity of GlmR is also critical to supplement UDP-GlcNAc synthesis. We also reveal that UDP-GlcNAc binding inhibits the catalytic activity of GlmR. While lysine acetylation (Ac) promotes this inhibition, phosphorylation appears to weaken UDP-GlcNAc binding and thus may prevent self-inhibition. **(B)** De novo amino sugar biosynthetic pathway involved in UDP-GlcNAc production. We predict that GlmM inhibition of CdaA is moderate when the forward reaction is effective such as when GlmR is present. In the absence of GlmR, we speculate carbon flow in this pathway is decreased where CdaA is not quite inhibited by GlmM which in turn leads to increased CdaA activity and elevated intracellular c-di-AMP. GlmU has two redox-sensitive cysteines that negatively affect its acetyltransferase activity in non-reducing conditions. **(C)** Speculative model depicting the main UDP-GlcNAc utilization pathways. The actin-like proteins MreB, MreBH, and Mbl assist with RodA/class B PBP machinery; PBP1 is a class A bifunctional enzyme. MreB/MreBH control cell width. Mbl is believed to monitor cell elongation. PBP1 is considered a general-purpose cell wall repair enzyme. Proper deployment of these four pathways is critical for cell width regulation. **(D)** In the absence of GlmR, we predict less accumulation of UDP-GlcNAc. We believe that this specifically impairs MreB pathway. **(E)** We suspect that overproduction of GlmR leads to elevated UDP-GlcNAc level which is specifically consumed through the MreB pathway. Hence, the decrease in cell width we observe in cells overexpressing *glmR* (Fig. 3). Arrows: dashed and increased thickness are used to indicate weak and strong flow/consumption respectively.

## DISCUSSION

### Antibiotic resistance

The enzymatic role of GlmR in drawing carbon for UDP-GlcNAc synthesis is now evident (**Fig. 8AB**). As such, cells devoid of *glmR* are incapable of mounting a proper response to host immune factors and cell wall targeting antibiotics (7, 8, 27, 72–74). Given that UDP-GlcNAc is also an essential component of WTA pathway, we show that cells lacking *glmR* are highly sensitive to tunicamycin (**Fig. 2**), a dual-function antibiotic that preferentially targets WTA at lower concentrations and PG biosynthesis at higher concentrations.

### Cell morphogenesis

The push and pull of PBP1 and MreB activities determine the cell width of *B. subtilis* cells (17–19). The similarities between *glmR* and *mreB* phenotypes hinted at an analogous role for GlmR (**Fig. 1C**). We now confirm that GlmR is a key cell width determinant (**Fig. 3E**). Besides MreB, *B. subtilis* encodes two additional MreB-like proteins Mbl and MreBH (**Fig. 8C** and **Fig. S6A**) (16, 75). Although *mreB* and *mbl* are essential and *mreBH* is not, the essentiality could be reversed in certain growth or genetic conditions (75). While MreB and MreBH play a role in the same pathway to achieve cell width control, Mbl is involved in an independent but parallel pathway to facilitate cell elongation (76, 77). In a high-throughput CRISPR-based analysis, it was recently found that *mbl* can be rendered non-essential in the absence of *glmR* (78). This is in direct contrast to MreB, where *glmR* overexpression supports the viability of cells lacking *mreB* (**Fig. 1C**) (15). Thus, it may be reasoned that GlmR differentially modulates MreB, Mbl, and PBP1 functions. However, based on the enzymatic function of GlmR, we speculate that the differences in synthetic essentiality are likely due to the different rate of PG precursor consumption and/or PG hydrolysis in these mutants, which can be supported by either *glmR* deletion (for Δ*mbl*) or overexpression (for Δ*mreB*). In the absence of GlmR, we expect a reduction in UDP-GlcNAc level (**Fig. 8B**), which disproportionately affects MreB (**Fig. 8D**). This is supported by the finding that only overexpression of *mreB* restores the cell morphology defects of cells lacking *glmR* - but not *mbl* or *mreBH* (15). Upon GlmR overproduction, we predict that UDP-GlcNAc level would be elevated and specifically favors MreB pathway, thus leading to decreased cell width (**Fig. 8E**).

Another genetic condition that leads to Δ*glmR* cell shape restoration is deletion of *pbp1* (**Fig. 1C**) (15). Eliminating the consumption of UDP-GlcNAc by PBP1 likely frees up this precursor for MreB. Additionally, *pbp1* deletion triggers alternative sigma factor SigI activation and consequently upregulation of *mreBH* ensues (**Fig. S6AB**) (79). This likely reinforces UDP-GlcNAc utilization by the collaborative MreB/MreBH pathway (76). This may also explain why deletion of *pbp1* or SigI activation suppresses the aberrant cell shape defects of both *mreB* and *mbl* knockout strains (75, 80). As depicted in **Fig. S6A**, the morphology of cells lacking *mbl* is quite distinct from Δ*mreB* mutant (81–83). In the extended discussion (see supplemental file), we use our working model to explain this peculiar Δ*mbl* phenotype, why deletion of *glmR* renders *mbl* non-essential, and why glucose and magnesium supplementation fail to restore the cell width of Δ*glmR* cells to resemble WT (**Fig. 3E**). Overall, we believe that GlmR dictates the cell width by supplying UDP-GlcNAc to bolster MreB activity. We do note that our rather simplistic model does not take other factors with known role in cell shape regulation into account (77, 84–87). Thus, further model refinement is warranted when more information regarding GlmR comes to light.

### Cytokinesis

We observed that the positioning of cytokinetic machinery is impaired when GlmR is absent (**Fig. 3B**). The underlying mechanism for this is unclear at the moment.

Specifically, how division site positioning systems are bypassed needs to be investigated. It is possible that altered carbon metabolism, turgor pressure, cell morphology, and/or membrane fluidity may lead to aberrant cytokinesis (88–92). Alternatively, it could be due to decreased WTA composition as cells lacking *tagO* also exhibit aberrant septation (93).

### Connections to c-di-AMP

In *B. subtilis* and other organisms, GlmM inhibits CdaA (**Fig. 8B**) (29–33). GlmR increases the substrate availability for GlmM by stimulating GlmS. GlmR also contributes to the removal of the product made by GlmM (GlcN1P) to generate UDP-GlcNAc. Thus, it is conceivable that both actions of GlmR would promote the enzymatic function of GlmM to favor the forward reaction. We suspect that this may in turn result in moderate inhibition of CdaA activity, which allows WT cells to maintain a relatively high intracellular c-di-AMP concentration (**Fig. 4**). Conversely, absence of GlmR would result in GlmM idling longer (or performing the reverse reaction) which may weaken the ability of GlmM to inhibit CdaA (**Fig. 8B**) and consequently promote increased c-di-AMP synthase activity. Thus, this nucleotide messenger may be used by cells to broadcast the status of GlmM function. Instead (or in addition), changes in turgor pressure due to a weakened cell wall (61), which we observe in Δ*glmR* strain manifested by abnormal cell morphology, could lead to increased enzymatic activity of CdaA. Therefore, the cytoplasmic regulation of CdaA by GlmM combined with the extra-cytoplasmic regulation of CdaA by CdaR (**Fig. 4C**), both detecting flawed cell wall synthesis in the absence of GlmR function may result in elevated c-di-AMP level (55, 61, 94). However, this reasoning fails to explain why we continue to see elevated c-di-AMP levels in the absence of GlmR even with glucose or magnesium supplementation (**Fig. 4B**).

Importantly, the aberrant cell morphology of Δ*glmR* strain is corrected with chemical supplementation. Therefore, an additional pathway may work in tandem with the above-mentioned possibilities, for example triggered by less-active MreB (**Fig. 8D**).

Alternatively, several mutations that suppress the deleterious Δ*glmR* phenotypes map to sites that permanently increase the transcription of *cdaA*-*cdaR*-*glmM*-*glmS* genes (8, 65). As such, there may be a compensatory regulatory mechanism that leads to upregulation of these genes when GlmR is absent. Therefore, increased level of CdaA could lead to elevated intracellular c-di-AMP concentration in cells lacking *glmR* regardless of glucose or magnesium supplementation. We find that increased expression of *glmM* in Δ*glmR* is sufficient to support WT-like growth and cell morphology even when *cdaA* is deleted (**Fig. S2C**). Thus, we can postulate that the reduction of intracellular c-di-AMP level is not deleterious by itself and may signify when the cell envelope is in need of reinforcement such as when the PG precursor pathway is inhibited or in the presence of cell wall stressors.

One of the main functions of c-di-AMP is to regulate potassium influx/efflux for osmoregulation (14, 95–103). We elaborate on the possible implications in the extended discussion provided in the supplemental file. Additionally, secreted c-di-AMP (63) and/or stringent response activation (104–108) may influence metabolic pathways in the absence of GlmR. Thus, further experiments are needed to test these possibilities.

### Enzymatic activity

GlmR and its homologs possess uridyltransferase activity to synthesize UDP-GlcNAc (**Fig. 8A**) (27). We show that the catalytic function of GlmR is: (i) essential for growth on DS, (ii) required to support WT-like growth on LA, and (iii) necessary to correct Δ*glmR* cell morphology defects (**Fig. 5**). The *B. subtilis* GlmU (GcaD; an essential protein) has both acetyltransferase (109) and uridyltransferase (110) functions to generate N-acetyl glucosamine-1-phosphate (GlcNAc-1P) and UDP-GlcNAc respectively (**Fig. 1A** and **Fig. 8AB**). Based on our results, we can infer that on DS, the uridyltransferase activity of GlmU is disabled and cells exclusively depend on GlmR (**Fig. 5C**). Could GlmR perform both functions of GlmU? This possibility was ruled out in an in vitro experiment with purified *L. monocytogenes* GlmR (27). Whether in vivo acetylation of GlmR (**Fig. 7**) allows for it to be an acetyltransferase remains to be seen. The key lysine residue is mutated in both acetyl-mimetic (K296Q) and acetyl-ablative (K296R) mutants of GlmR, yet they complement Δ*glmR* phenotypes. Thus, perhaps one of the acetylated partners of GlmR such as YvcJ may serve as an acetyl donor. Intriguingly, the morphogenetic protein MreB (discussed above) is also acetylated at three different residues and the acetyl-ablative mutation of one of the lysines impairs cell morphology control (38).

Additional experiments are necessary to investigate these models.

In *Mycobacterium tuberculosis*, phosphorylation of GlmU by an S/T kinase was found to negatively affect the acetyltransferase activity but not uridyltransferase activity (111).

Thus, one of the two functions of GlmU could be specifically deactivated by post-translational modification. It was found that *glmU* gene expression is downregulated during stringent response (112, 113). The production of stringent response alarmone, (p)ppGpp, can be elicited by a c-di-AMP binding protein, DarB (106). As we observed increased c-di-AMP level in the absence of GlmR (**Fig. 4**), perhaps on DS medium *glmU* transcription is repressed by stringent response activation. Pyruvate kinase is another metabolic enzyme with link to stringent response (114), which is differentially autoregulated in glycolytic vs gluconeogenic conditions (115). Thus, it is possible that GlmU activity is affected similarly. Additionally, conditions such as high magnesium and low pH hinder the acetyltransferase activity of *B. subtilis* GlmU (109). Intriguingly, alteration in c-di-AMP levels is also known to modulate intracellular magnesium concentration (14). Moreover, it has been found that GlmU is regulated by two cysteine residues that are involved in disulfide bond formation and therefore responsive to intracellular redox status (**Fig. 8B**) (109, 116). Specifically, acetyltransferase activity was found to be diminished in non-reducing conditions. It is therefore plausible that the intracellular conditions of Δ*glmR* mutant in DS growth medium are not optimal for one or both enzymatic function(s) of GlmU. Intriguingly, UDP-GlcNAc is also essential for bacillithiol production (**Fig. 1A**). Bacillithiol is an important agent involved in redox stress response as well as a resistance factor for cell wall targeting antibiotics (117).

Therefore, it is tempting to speculate that in the absence of GlmR cells are unable to properly respond to oxidative stress. However, overexpression of *glmS*/*glmM* (**Fig. S2BC**) appears to fully restore GlmU activity perhaps by alleviating oxidative stress. Alternatively, the cause of death of Δ*glmR* cells and failure to complement by enzymatically inert mutant could be indirect. For example, this mutant may fail to stimulate GlmS and draw carbon for GlmU to execute its function (**Fig. 8B**).

Experiments to test these speculations are underway.

### Phosphorylation

Prior to the knowledge of its uridyltransferase activity, GlmR was shown to bind UDP-GlcNAc. The mutants defective in binding UDP-GlcNAc did not exhibit any noticeable phenotype (8, 34). However, we show that the binding of UDP-GlcNAc ligand moderates GlmR function (**Fig. 8A**) as overproduction of UDP-GlcNAc binding deficient variant (R301A) is toxic (**Fig. 6AB**). Similarly, phosphoregulation of GlmR and its physiological importance have been investigated previously (7, 8). GlmR mutants either mimicking phosphorylated or unphosphorylated state support growth. PrkC, the kinase responsible for phosphorylating GlmR, is known to sense cell wall stress and/or the changes in the levels of PG precursors and finetune growth rate accordingly (35–37, 118). Thus, it is possible that phosphorylation of GlmR may serve to prioritize carbon for strengthening the cell envelope through boosting UDP-GlcNAc level. Subsequently, the removal of phosphate by PrpC phosphatase may lower the function back to basal level but not completely off - this would explain why phosphoablative mutant is also able to complement. Results of our in vitro experiments suggest that phosphorylation may hinder but not abolish UDP-GlcNAc binding (**Fig. 8A**). Intriguingly, while overexpression of the mutant deficient in recognizing UDP-GlcNAc (R301A) is toxic, phosphomimetic mutant (T304E) where UDP-GlcNAc binding appears to be more transient is not.

However, GlmR is subject to multiple levels of regulation within the cell that cannot be incorporated with purified proteins. As only phosphomimetic version of GlmR supports rod shape maintenance in Δ*mreB* (7), we could envision a scenario where phosphorylation increases UDP-GlcNAc synthesis and support Mbl/MreBH pathway to counteract the action of PBP1 (**Fig. 8C**). On the contrary, phosphoablative mutant is unable to sufficiently elevate the UDP-GlcNAc level in Δ*mreB* strain background to support rod shape maintenance. Alternatively, as MreB and PBP1 form a complex (17) and GlmR assists in PBP1 localization (7, 15), perhaps GlmR phosphorylation alters the partner preference. Further investigation is required to test these hypotheses.

### Acetylation

Similar to phosphorylation, acetylation of lysine residues serves as another reversible post-translational regulatory mechanism (71). A handful of proteomics studies have identified acetylated proteins in *B. subtilis* (38, 39, 69, 70). GlmR is found to be acetylated along with several other proteins including GlmS, GlmM, GlmU, and YvcJ discussed in this report (**Fig. 1A**). Therefore, we investigated whether acetylation influences GlmR function. Our results revealed that GlmR acetyl-mimetic (K296Q) mutant was equally efficient as WT copy in complementing *glmR* phenotypes. However, acetyl-ablative GlmR mutant (K296R) was toxic upon overexpression similar to the mutant (R301A) that is unable to bind UDP-GlcNAc. Based on our results, it appears that acetylation of lysine 296 of GlmR may serve to negatively regulate its enzymatic activity (**Fig. 8A**). Interestingly, the equivalent residue is naturally glutamine in *Bacillus halodurans* and based on the crystal structure (PDB ID: 2O2Z) it does not appear to be involved in the coordination of its ligand, NAD in this case (34). However, this loop-like region is not clearly resolved in the crystal structure and is possibly dynamic. In *Staphylococcus aureus* and *Listeria monocytogenes*, the corresponding residue appears to be aspartate and glutamate respectively (27). Nonetheless, other proximal lysine residues are present and they may be acetylated. Therefore, acetylation is likely an additional post-translational mechanism that finetunes the function of GlmR (119–121). Besides acetylation, GlmR appears to be also subject to succinylation and malonylation (70). Thus, multiple modes of regulation may exist.

### Conservation of GlmR and its potential as drug target

Our findings about *B. subtilis* GlmR are likely broadly applicable to its homologs in other organisms which include several clinically relevant pathogens. GlmR is highly conserved in diverse bacterial lineages and in some archaea (9, 26). However, except for Gram-positive bacteria not much is known about the function of GlmR-like proteins in other organisms. In *M. tuberculosis* (CuvA/Rv1422), it is found to accumulate at sites of active cell wall synthesis and contribute to virulence (72). It is also found to be phosphorylated by an S/T kinase in this organism (122). *L. monocytogenes* GlmR is important for survival within the host during infection, and it is also phosphorylated by an S/T kinase (73). In *S. aureus*, *glmR* is an essential gene (123, 124). However, another study indicates that *glmR* mutant is temperature sensitive (125). A link between GlmR, S/T kinase, and CdaA appears to exist in *S. aureus* as well (126). *Enterococcus faecalis* GlmR phenotype differs from that of its *B. subtilis* counterpart (74). Nevertheless, it binds to UDP-GlcNAc and plays a role in antibiotic resistance. GlmR homologs of the above-mentioned organisms are enzymatically active and produce UDP-GlcNAc (27, 74). The specific function of *E. coli* homolog (YbhK) is yet to be characterized. However, *yhbK* of *E. coli* was found to complement *glmR* in *B. subtilis* – thus it may play a similar enzymatic function in its host organism as well (9). Disruption of *yhbK* is associated with increased tolerance to the cell wall targeting antibiotic ampicillin (127). Intriguingly, GlmR in *Salmonella enterica* is regulated via glycosylation, specifically with GlcNAc addition to an arginine (128). Thus, additional experiments are needed to uncover the full scope of the regulatory modes of GlmR and other related pathways. Given its crucial role in antibiotic resistance and pathogenesis, inhibitors of GlmR enzyme could be developed as novel therapeutics.

### Summary

*B. subtilis* is an excellent model to investigate fundamental biological pathways.

Additionally, it is also a widely used bacterium in various biotechnology industry (1, 129). Thus, deeper understanding of metabolic processes would propel the fields of both basic and applied biology. In this report, we show that GlmR is a central metabolic enzyme which is regulated by multiple means. More specifically our results suggest that post-translational regulation of GlmR may aid in decisions involving carbon prioritization. This role becomes vital in times of cell envelope stress. Therefore, unsurprisingly, cells lacking *glmR* are highly susceptible to cell envelope targeting antibiotics. As antibiotic resistance is a growing concern worldwide, key metabolic pathways and proteins such as GlmR could serve as potential therapeutic targets (116, 130–133).

## MATERIALS AND METHODS

### Strain construction

*B. subtilis* strains utilized in this study are derivatives of PY79 (134). Specific details regarding strain construction can be found in the supplemental file. Table S1 contains all strains and oligonucleotides referenced. The knockout strains were obtained from the Bacillus Genetic Stock Center (135). Plasmids pDG1662 (136), pDR111 (David Rudner), pDR244 (135, 137), and pBS2EXylRP*_xylA_* (ECE741; (138)) were used to create strains for this study. pET28a vector was used for purifying recombinant *B. subtilis* GlmR and its mutants from *E. coli*. QuikChange kit (Agilent) was used for site-directed mutagenesis. All plasmids generated were verified through Sanger or whole plasmid sequencing (Azenta). All *B. subtilis* chromosomal gene deletions and insertions were confirmed via PCR and other standard techniques.

### Media used and spot titer assay

Overnight cultures grown at 30 °C in lysogeny broth (LB) were serially diluted up to 10^-5^ and 1 µL of culture was plated on LB agar (LA) or BD Difco Starch agar (DS) plates containing either 0 or 1 mM IPTG. Plates were imaged after overnight incubation at 37°C.

### Antibiotic susceptibility testing

Zone of inhibition (ZOI) assays were completed after overnight cultures of the indicated *B. subtilis* strains were grown in LB at 30 °C. Cultures were standardized to an OD_600_ of 0.1 and 100 µl was spread using sterile glass beads on LA plates. Plates were allowed to dry with their lid off for 30 minutes in a biosafety cabinet. Plates with sterile filter paper discs laced with 5 µl of 0, 10, 25, 50, or 100 µg/ml tunicamycin were incubated overnight at 37 °C. ZOI diameter quantification was performed via FIJI (139). The standard diameter of the filter paper discs (6.5 mm) was subtracted from all ZOI measurements.

### Western blot

Immunoblot analysis of indicated strains were completed after overnight cultures of *B. subtilis* strains grown in LB at 30 °C were diluted to an OD_600_ of 0.05 in 10 ml of LB. When indicated, cultures were supplemented with a final concentration of 25 mM MgCl2, 1% glucose, or 1 mM IPTG. Cultures were grown to an OD_600_ of ∼1, standardized to an OD_600_ of exactly 1 before centrifugation and resuspension in protoplast buffer containing 0.5 M sucrose, 20 mM MgCl_2_, 10 mM KH_2_PO_4_, and 0.1 mg/ml lysozyme. Samples were incubated at 37 °C for 30 min and then prepared for SDS-PAGE. After electrophoresis, the samples were transferred onto a nitrocellulose membrane and subsequently probed with appropriate antibodies.

### Microscopy

Overnight cultures of *B. subtilis* strains grown at 30 °C in LB were diluted to an OD_600_ of 0.05 in 10 ml LB. If noted, cultures were made to a final concentration of 25 mM MgCl_2_, 1% glucose, or 1 mM IPTG. For the time course microscopy, cultures were grown for the duration indicated. For all other experiments, cultures were grown to an OD_600_ of 1 at 37 °C. Sample preparation and microscopy techniques were completed as previously described (140).

### Quantification and statistics

Analysis of ZOI, colony size, and measurements of cell length/width were performed using FIJI (139). Statistical analysis was conducted using GraphPad Prism version 10.4.1. Ordinary one-way ANOVA with multiple comparisons was used with Tukey’s method.

### Biochemical analysis

For labeling WT and mutant forms of GlmR, the purified proteins were labeled with fluorescein isothiocyanate (FITC) as previously reported (68). Briefly, 50 µM of each WT-GlmR and mutants were incubated with FITC (250 µM) in 50 mM phosphate buffer (pH 8.0) for 5 h on ice. The reaction was stopped by adding 5 mM Tris-HCl (pH 8.0).

Sephadex G25 fine column (Cytiva) was used to separate the free FITC from the labeled proteins. The concentrations of FITC-bound GlmR and mutants were determined by absorbance at 495 nm. The concentrations of the proteins were measured using Bradford assay. The stoichiometry of labeling FITC per GlmR and mutant constructs was found to be approximately 0.6.

## ACKNOWLEDGEMENTS

We thank the members of our laboratory for comments on the manuscript. This work was funded by the National Institutes of Health grant R35GM133617 and the University of South Florida - Center for Antimicrobial Resistance award (P.J.E.).

## AUTHOR CONTRIBUTIONS

Study design (L.S., S.K., D.B., and P.J.E.), strain construction and data acquisition (L.S., S.K., D.B., and S.D.), data analysis (L.S., S.K., D.B., S.D. and P.J.E.), and writing of the manuscript (L.S., S.K., D.B., and P.J.E.).

## Notes

### Competing Interest Statement

The authors have declared no competing interest.

