## Supplemental Text and Figures for "Staying in the loop to make ends meet: roles and regulation of GlmR in *Bacillus subtilis*"

### Supplemental Results and Discussion

*Rescue of ΔglmR phenotypes by additional deletion of cdaA is likely due to polar effect*

Our data reveal that the intracellular concentration of c-di-AMP is elevated in Δ*glmR* cells through a pathway that requires CdaA and CdaR (**Fig. 4**). Therefore, to investigate whether alteration in c-di-AMP concentration affects Δ*glmR* cell morphology we used fluorescence microscopy (**Fig. S2**). For this purpose, we examined the cell morphologies of strains lacking *disA*, *cdaA*, and *cdaR* individually and in combination with *glmR* (**Fig. S2A**). The cell morphologies of Δ*disA*, Δ*cdaA*, and Δ*cdaR* strains resembled WT. Therefore, depletion of intracellular c-di-AMP concentration alone does not affect cell morphology in our growth conditions. Conversely, Δ*disA* Δ*glmR* double-deletion mutant appeared similar to Δ*glmR*. Remarkably, in the Δ*cdaA* Δ*glmR* and Δ*cdaR* Δ*glmR* strains the typical abnormal Δ*glmR* cell morphology was completely abrogated. However, this result is counterintuitive as previous reports show only conditions that elevate c-di-AMP level, such as deletion of c-di-AMP phosphodiesterases alleviate the severity of Δ*glmR* phenotypes (1, 2). Therefore, we wondered if the reversal of Δ*glmR* phenotype could be due to the polar effect stemming from the antibiotic cassette mediated disruption of *cdaA* or *cdaR* (**Fig. S2B**) - which introduces an additional promoter to drive downstream gene expression (3). More specifically, as GlmM is encoded from the highly conserved *cdaA*-*cdaR*-*glmM* operon (4), we suspected that the abrogation of Δ*glmR* phenotype by deletion of *cdaA* or *cdaR* could be explained by increased expression of *glmM* (**Fig. 1A**). Overproduction of GlmM has been shown to rescue Δ*glmR* phenotypes previously (2, 5). Additionally, mutations that suppress Δ*glmR* phenotypes map to the *cdaA* locus and result in increased expression of *glmM* and/or *glmS* (**Fig. S2B**; red asterisks) (2, 6). Therefore, we tested our prediction through two independent approaches - by either removing the antibiotic resistance marker from Δ*cdaA* (Δ*cdaA**) or by overexpressing *glmM* (*glmM*$^+$) from an ectopic locus in the Δ*cdaA** background. As anticipated, unlike Δ*cdaA* Δ*glmR*, deletion of *glmR* in markerless Δ*cdaA** background did not fully correct the abnormal Δ*glmR* cell morphology (**Fig. S2C**). On the contrary, increased expression of *glmM* in Δ*cdaA** Δ*glmR* background was sufficient to restore WT-like cell morphology.

We also employed the plate assay described in the main text to investigate the polar effect further (**Fig. 5**). In **Fig. S3A**, we show that the Δ*glmR* growth phenotypes (poor growth on LA and severe growth inhibition on DS) is corrected by the synthetic expression of inducible *glmS* from an ectopic locus. This effect was seen in MH medium as well (2). Next, we monitored the growth characteristics of Δ*cdaA* Δ*glmR* and Δ*cdaA** Δ*glmR* double mutants. Intriguingly, both strains grew better than *glmR* single deletion on LA and DS plates (**Fig. S3B**). However, the growth of the former appears to be better than the latter. On DS, the growth of markerless version (Δ*cdaA** Δ*glmR*) was also significantly (>4-log) enhanced when compared to the Δ*glmR* strain. This is possibly due to the altered proximity to the native promoter which presumably leads to increased *glmM* expression (**Fig. S2B**). We also ensured that both Δ*cdaA* and Δ*cdaA** do not exhibit any growth defect on their own (**Fig. S3C**). Introduction of inducible *glmM* in Δ*cdaA** Δ*glmR* strain promoted growth on LA and DS even in the absence of inducer due to leaky expression. However, in the presence of inducer, the growth characteristics resembled that of WT on both plates (**Fig. S3B**). Therefore, our results indicate that the reversal of Δ*glmR* cell morphology and growth phenotypes by *cdaA* deletion could be explained by the polar effect stemming from enhanced expression of *glmM*. Thus, researchers should exercise caution in interpreting results with knockout strains – even those with antibiotic resistance cassette removed.

*Deletion of gdpP, pgpH, or disA enhances the growth of ΔglmR mutant*

To investigate the possible effect of other c-di-AMP related proteins (**Fig. 4A**), we tested the growth of the Δ*glmR* strain harboring an additional deletion of a phosphodiesterase gene of either GdpP or PgpH. First, we confirmed that all the individual knockout strains grow similar to WT on both media types (**Fig. S3C**). On LA, we find that the *glmR gdpP* and *glmR pgpH* double deletion strains resemble Δ*glmR* (**Fig. S3D**). However, on DS, we see significantly enhanced growth for Δ*glmR* Δ*gdpP* and mild improvement for Δ*glmR* Δ*pgpH*. This is consistent with previous findings (1, 2). It is to be noted that the growth of these strains are not quite similar to WT based on the opacity of the colonies on DS medium. We also investigated the growth of *glmR disA* double mutant (**Fig. S3E**). The growth of strain harboring *disA* single deletion resembled WT control on both LA and DS. On LA, the growth phenotype of the Δ*disA* Δ*glmR* double deletion mutant appeared more similar to Δ*glmR*. Surprisingly, on DS, we find that this strain lacking DisA and GlmR grows nearly 4-log better than the Δ*glmR* single deletion strain. Thus, it appears that the absence of DisA is beneficial for the growth of Δ*glmR* cells in this specific experimental condition.

Why both decrease (Δ*disA*) and increase (Δ*gdpP/*Δ*pgpH*) of c-di-AMP levels promote the growth of cells lacking *glmR* is puzzling (**Fig. S3DE**). One possibility is that perhaps when *disA* and *glmR* are deleted, cells compensate with increased expression of *cdaA*, and thus *glmM* as well

(**Fig. S2B** and **Fig. 4C**). This may mildly alleviate the growth phenotype of Δ*disA* Δ*glmR*. Altered production of c-di-AMP synthases in the absence of other c-di-AMP enzymes have been noted before in *B. subtilis* (7, 8). However, this prediction needs to be tested. Regarding phosphodiesterases, previous observations already demonstrated that disruption of *gdpP* or *pgpH* alleviates *glmR* deletion phenotypes (1, 2). Absence of GdpP or PgpH would likely elevate the intracellular c-di-AMP concentration (7, 9). Given that the level of c-di-AMP may already be higher in Δ*glmR* cells (**Fig. 4C**), additional deletion of *gdpP*/*pgpH* would possibly further elevate the intracellular c-di-AMP concentration. Thus, it can be presumed that the additional increase in c-di-AMP level is critical for promoting the growth of cells lacking *glmR* specifically on DS (**Fig. S3E**). It is to be noted that although the growth phenotype on plate is somewhat alleviated, the aberrant Δ*glmR* cell morphology is not corrected by *gdpP* or *pgpH* deletion (**Fig. S2D**). High intracellular c-di-AMP concentration is known to assist with cell wall stress response (9, 10). However, the precise mechanism as to why deletion of one of the phosphodiesterases in cells lacking *glmR* supports growth remains unclear. Our data suggests that GlmU function may become impaired on DS (**Fig. 5C**). Thus, we can speculate that the deletion of *gdpP*/*pgpH* in Δ*glmR* supports GlmU to remain at least partially active. Or perhaps elevated c-di-AMP level may alter the intracellular magnesium concentration (11) and/or enhance the production or function of GlmS (**Fig. S3A**) to subsequently bolster GlmU activity. Additional experiments are warranted to test these hypotheses.

*Role of GlmR in cell morphogenesis*

In our model, we propose that the balanced UDP-GlcNAc utilization by the MreB, Mbl, MreBH, and PBP1 pathways allows for cell width maintenance in WT (**Fig. S6A**). There is evidence to suggest that MreB and MreBH are participating in the same pathway while Mbl works independently (12-14). It has been reported that the phenotypes of *mreB*/*mbl* overexpression are different (15). While MreB or MreBH overproduction is toxic, Mbl is not – thus hinting that

Mbl function is possibly moderated in the presence of MreB/MreBH/PBP1 (thus the arrow thickness for Mbl remains unchanged in all the panels of **Fig. S6** and **Fig. 8CDE**). As per our model, in the absence of *mreB*, UDP-GlcNAc is consumed predominantly by PBP1 which causes cell bulging (**Fig. S6A**). This effect can be relieved by *glmR* overexpression to elevate the supply of UDP-GlcNAc to support the functions of Mbl and MreBH (**Fig. 1C**). Alternatively, deletion of *pbp1* also corrects the cell shape defects of *mreB* deletion strain through SigI-mediated upregulation of *mreBH* (16). However, according to our model, deletion of *pbp1* alone would strongly favor the MreB/MreBH pathway and result in decreased cell diameter (**Fig. S6A**). We suspect that the absence of Mbl promotes increased UDP-GlcNAc usage by PBP1 and MreB pathways giving rise to its unique spiral cell morphology (**Fig. S6A**). As such, the rescue of Δ*mbl* viability can be achieved by additional deletion of *glmR* (14), as this would lead to less active MreB as well as decreased UDP-GlcNAc accumulation to limit PBP1 consumption.

Our results show that neither glucose nor magnesium supplementation fully restores the cell width of Δ*glmR* cells similar to WT (**Fig. 3E**). We suspect that without GlmR, the tightly linked GlmR/MreB pathway becomes dysfunctional and UDP-GlcNAc made by GlmU (in the presence of glucose or magnesium) is primarily utilized by PBP1 and Mbl which supports rod shape maintenance. However, in the absence of GlmR, MreB is unable to fully participate. Thus, although rod shape is restored, cell width remains larger than WT in Δ*glmR* cells grown in the presence of glucose or magnesium.

*Potential role for potassium:*

Differential inhibition of potassium channels by low vs high c-di-AMP concentrations has been noted (17). After osmotic upshift, c-di-AMP level drops immediately to favor intracellular potassium accumulation (and concomitant magnesium export) temporarily (11). It is known that potassium response to osmotic stress is more transient, and alternative compatible solutes such as glycine betaine and proline are subsequently favored (18). Therefore, perhaps altered potassium level and subsequent dysregulated osmotic stress response activation could presumably be the source of Δ*glmR* toxicity. Intriguingly, polymerization of MreB is inhibited by high intracellular potassium concentration (19). Elevated levels of c-di-AMP would limit potassium influx and promote extrusion (20), which may therefore support MreB polymerization. However, as discussed above MreB is likely less active without GlmR – therefore, the predicted increase in c-di-AMP level in *glmR gdpP/pgpH* double mutants are unable to correct the abnormal cell morphology of Δ*glmR* (**Fig. S2D**). Intriguingly, high potassium concentration also dramatically alters the polymerization kinetics of the tubulin-like protein FtsZ, which determines the cell division site (21). Therefore, it is possible altered potassium level may contribute to the division site positioning defect that we observe in cells lacking *glmR* (**Fig. 3B**). In addition to its role in osmotic stress response, potassium is also important for the maintenance of intracellular pH, membrane potential, ion homeostasis, and ribosome function to name a few (22-24). As such, additional experiments are warranted to elucidate the different pathways and possibilities.

__Supplemental Figures and Corresponding Figure Legends__

**Supplementary figure 1: GlmR is important for correct cell shape and septation.** Representative micrographs of WT (PY79) and Δ*glmR* (RB176) strains grown in LB in the absence or presence of D-glucose (1%) or magnesium (25 mM $MgCl_2$) supplementation, imaged hourly for four hours. Yellow arrows indicate examples of abnormal septation.

**Supplementary figure 2: Deletion of *cdaA* rescues Δ*glmR* phenotype likely due to polar effect. (A)** Fluorescence micrographs of membrane-stained (FM 4-64, red): WT (PY79), Δ*glmR* (RB176), Δ*disA* (SK97), Δ*disA* Δ*glmR* (SK102), Δ*cdaA* (SK97), Δ*cdaA* Δ*glmR* (SK101) Δ*cdaR* (SK130), or Δ*cdaR* Δ*glmR* (SK131). Scale bar, 1 µm. **(B)** Genetic locus of *cdaA*-*cdaR*-*glmM* operon. Genes *sigW-rsiW* are located upstream of this operon while *glmS* is present immediately downstream in the *B. subtilis* genome. Red asterisks indicate the position of mutations commonly found in the Δ*glmR* suppressors that allow increased transcription of *cdaA*-*cdaR*-*glmM* genes and/or *glmS*. The terminator downstream of this *cdaA* operon is weaker (depicted with shorter symbol) resulting in read-through transcription of *glmS*. *glmS* has its own promoter followed by a riboswitch-ribozyme (6, 25). Replacement of *cdaA* or *cdaR* with an antibiotic resistance cassette introduces additional promoter (shown in red) and removal of the cassette replacing *cdaA* (Δ*cdaA**; markerless) brings *glmM* closer to its native promoter, thus potentially result in stronger expression. **(C)** Fluorescence micrographs of membrane-stained (FM 4-64, red): WT (PY79), Δ*glmR* (RB176), Δ*glmR* Δ*cdaA** (SK138), and Δ*glmR* Δ*cdaA** with inducible *glmM*$^+$ (BLS67). When indicated, 0 and 1 mM IPTG was used in IPTG (-) and (+) conditions respectively. Scale bar, 1 µm. **(D)** Representative micrographs of WT (PY79), Δ*glmR* (RB176), Δ*glmR* Δ*gdpP* (SK103), and Δ*glmR* Δ*pgpH* (SK104). Red, FM 4-64 membrane stain. Scale bar, 1 µm.

**Supplementary figure 3: Single deletions of proteins related to c-di-AMP. (A)** Serial dilutions of WT (PY79), Δ*glmR* (SK35), Δ*glmR glmS*$^+$ (BLS84) on LA, LA + 3% xylose, DS, or DS + 3% xylose. **(B)** Serial dilutions of WT (PY79), Δ*glmR* (RB176), Δ*glmR* Δ*cdaA* (SK101), Δ*glmR* Δ*cdaA** (SK138), and Δ*glmR* Δ*cdaA* glmM*$^+$ (BLS67) on LA, LA + 1 mM IPTG, DS, or DS + 1 mM IPTG. **(C)** Spot titer assay of WT (PY79), Δ*glmR* (SK35), Δ*cdaA* (SK96), Δ*cdaA** (SK137), Δ*gdpP* (SK98), and Δ*pgpH* (SK99) on LA and DS. **(D)** Growth of serially-diluted culture aliquots of WT (PY79), Δ*glmR* (RB176), Δ*glmR* Δ*gdpP* (SK103), and Δ*glmR* Δ*pgpH* (SK104) on LA and DS plates. **(E)** Spot titer assay showing growth of WT (PY79), Δ*glmR* (RB176), Δ*disA* (SK97), and Δ*glmR* Δ*disA* (SK102) on LA and DS plates.

**Supplementary figure 4: GlmR D38A D39A is non-functional and stably produced**. **(A)** Serial dilutions of WT (PY79), Δ*glmR* (SK35), Δ*glmR glmR*$^+$ (SK56), Δ*glmR glmR-6his*$^+$ (BLS101), and Δ*glmR glmR-D38A-D39A-6his*$^+$ (BLS102) on LA, LA + 1 mM IPTG, DS, or DS + 1 mM IPTG. **(B)** Representative western blot of Δ*glmR glmR-6his*$^+$ (BLS101) and Δ*glmR glmR-D38A-D39A-6his*$^+$ (BLS102) with IPTG (1 mM) induction probed with anti-His antibody. Ponceau S-stained total protein gel for both samples serves as loading control.

**Supplementary figure 5**: **Phosphomutants of GlmR are functionally similar to WT. (A)** Spot titer assay of WT (PY79), Δ*glmR* (SK35), Δ*glmR glmR*$^+$ (SK56), Δ*glmR glmR-T304A*$^+$ (SK139), and Δ*glmR glmR-T304E*$^+$ (SK140) on LA, LA + 1 mM IPTG, DS, and DS + 1 mM IPTG. **(B)** Micrographs of WT (PY79), Δ*glmR* (SK35), Δ*glmR glmR*$^+$ (SK56), Δ*glmR glmR-T304A*$^+$ (SK139), and Δ*glmR glmR-T304E*$^+$ (SK140) with or without 1 mM IPTG induction. **(C)** Cell width quantifications of Δ*glmR glmR*$^+$ (SK56), Δ*glmR glmR-T304A*$^+$ (SK139), and Δ*glmR glmR-T304E*$^+$ (SK140). **(D)** Colony area measurements relative to uninduced Δ*glmR glmR*$^+$ (SK56), Δ*glmR glmR-T304A*$^+$ (SK139), or Δ*glmR glmR-T304E*$^+$ (SK140). Area measured via FIJI automatically using thresholding. One-way ANOVA with Tukey's correction was used for interpreting statistical significance; * = $p<0.05$, ** = $p<0.01$, ns = $p>0.05$.

**Figure S6: Addendum to the working model. (A)** As described in Fig. 8, UDP-GlcNAc produced by GlmR and GlmU enzymes is consumed through the pathways involving MreB, MreBH, Mbl, and PBP1 in WT cells. In the absence of MreB, hyperactive PBP1 leads to abnormal cell bulging. This consequence is averted by either deletion of *pbp1* or overexpression of *glmR* (depicted in Fig. 1C). When PBP1 is absent, alternative sigma factor SigI is activated which in turn upregulates *mreBH*. Therefore, the combined action of MreB and MreBH involved in cell width control leads to decreased cell width. In cells lacking *mbl*, UDP-GlcNAc utilization happens through both MreB and PBP1 pathways which result in twisted cell morphology. Thus, either deletion of *glmR* (lowers UDP GlcNAc level) or *pbp1* (increases MreBH activity) restores viability. **(B)** In cells lacking *glmR*, MreB pathway is weakened and PBP1 becomes hyperactivated. This leads to cell shape abnormality. Thus, either overexpression of *mreB* or deletion of *pbp1* results in cell morphology correction. Weak and strong UDP-GlcNAc consumption are represented with dashed and thicker arrows respectively.

**Supplementary video 1: Aberrant positioning of cytokinetic machinery in the absence of GlmR.** This video shows Δ*glmR* (SK35) cell undergoing abnormal septation. Red, FM 4-64 membrane dye. Movie created in FIJI.

### Supplemental Methods

**Strain construction:**

*Bacillus subtilis* knockout strains were requested from the Bacillus Genetic Stock Center (BGSC). Chromosomal DNA of the strains harboring gene deletions were transformed into PY79 and confirmed using PCR. Plasmid pDR244 was used to generate markerless strains. Chromosomal insertion of genes of interest from integration vectors (pDG1662 or pDR111) were confirmed using standard protocol. *Escherichia coli* DH5a strain was used for plasmid maintenance. *E. coli* BL21-DE3 strain was used for protein purification.

**Plasmid construction:**

1. **c-di-AMP reporter**: The promoter harboring c-di-AMP riboswitch of *kimA* was amplified with oSK44/oSK45 from *B. subtilis* 168 chromosomal DNA and *gfp* with oSK46/oSK47. The resulting fragments were digested with EcoRI/NheI and NheI/BamHI, respectively, and ligated into pDG1662 digested with EcoRI/BamHI to create the plasmid pSK13.

2. **GlmR/mutant purification**: *glmR* was amplified from the PY79 chromosome using primers oSK48/oSK49; the resulting fragment was then digested with BamHI and NheI and ligated into pET28a (BamHI/NheI) to generate pSK14. The plasmid pSK14 was used for site-directed mutagenesis (QuikChange; Agilent) using primer pairs oSK60/oSK61, oSK56/oSK57, and oSK58/oSK59 to create versions of *glmR* harboring the R301A (pSK18), T304A (pSK17), and T304E (pDB04) mutations respectively.

3. ***glmR/mutant* complemention**: *glmR* was amplified from the PY79 genomic DNA using primers oSK16/oSK17; the resulting fragment was then digested with HindIII and SalI and ligated into pDR111 (HindIII/SalI), an IPTG-inducible vector for integration into the *amyE* locus in *B. subtilis* chromosome to generate pSK7. Similarly, *glmR* from pET28a plasmids harboring mutations R301A (pSK18), T304A (pSK17), and T304E (pDB04) were amplified using oSK16/oSK17 and cloned into pDR111 to generate pSK26, pSK24, and pSK25 respectively. pSK7 was amplified using the QuikChange kit (Agilent) with primer pairs oSK68/oSK69 and oSK70/oSK71 to introduce K296Q (pSK22) and K296R (pSK23) mutations respectively. To engineer the enzymatically-inactive GlmR mutant (D38A D39A), pSK7 was amplified with primer pairs oLS119/oLS120 to generate pLS59 using the QuikChange kit (Agilent). To generate *glmR-his6* and D38A D39A-his6, *glmR* from either pSK7 or pLS59 was amplified with oLS144/oLS145. The resulting fragments were

digested with SalI/NheI and ligated into pDR111 resulting in plasmids pLS79 and pLS80, respectively.

4. ***Other glm proteins***: *glmM* and *glmS* genes were amplified from PY79 genomic DNA using primer pairs oSK86/oSK87 and oLS69/oLS70 respectively. *glmM* fragment was digested with SalI/NheI and cloned into pDR111 to make oLS29. *glmS* PCR product was digested with XbaI/PstI and ligated into pBS2EXylRP$_{xylA}$ (ECE741; (26)) to generate oLS35.

**Table S1: Strains used in this study**

| Strain | Genotype | Reference |
|---|---|---|
| PY79 | Wildtype *B. subtilis* | (27) |
| RB176 | *glmR::erm* | BKE34760 (BGSC) → PY79 |
| SK29 | *glmR::erm*; *amyE::*$P_{hyperspank}$*-glmR spec* | pSK7 → PY79 |
| SK35 | Δ*glmR* (markerless) | pDR244 → RB176 |
| SK56 | Δ*glmR*; *amyE::*$P_{hyperspank}$*-glmR spec* | pSK7 → SK35 |
| SK94 | *amyE::PkimA-gfp-cat* | pSK13 → PY79 |
| SK96 | *cdaA::kan* | BKK01750 (BGSC) → PY79 |
| SK97 | *disA::kan* | BKK00880 (BGSC) → PY79 |
| SK98 | *gdpP::kan* | BKK40510 (BGSC) → PY79 |
| SK99 | *pgpH::kan* | BKK25330 (BGSC) → PY79 |
| SK101 | *cdaA::kan*; *glmR::erm* | SK96 → RB176 |
| SK102 | *disA::kan*; *glmR::erm* | SK97 → RB176 |
| SK103 | *gdpP::kan*; *glmR::erm* | SK98 → RB176 |
| SK104 | *pgpH::kan*; *glmR::erm* | SK99 → RB176 |
| SK107 | *glmR::erm*; *amyE::*$P_{hyperspank}$*-glmR-K296Q spec* | pSK22 → RB176 |
| SK108 | *glmR::erm*; *amyE::*$P_{hyperspank}$*-glmR-K296R spec* | pSK23 → RB176 |
| SK113 | *glmR::erm*; *amyE::*$P_{kimA}$*-gfp cat* | SK94 → RB176 |
| SK109 | *disA::kan*; *amyE::*$P_{kimA}$*-gfp cat* | SK94 → SK97 |
| SK110 | *cdaA::kan*; *amyE::*$P_{kimA}$*-gfp cat* | SK94 → SK96 |
| SK111 | *disA::kan*; *glmR::erm*; *amyE::*$P_{kimA}$*-gfp cat* | SK94 → SK102 |
| SK112 | *cdaA::kan*; *glmR::erm*; *amyE::*$P_{kimA}$*-gfp cat* | SK94 → SK101 |
| SK118 | *cdaR::kan*; *amyE::*$P_{kimA}$*-gfp cat* | SK94 → SK130 |
| SK119 | *glmR::erm*; *cdaR::kan*; *amyE::*$P_{kimA}$*-gfp cat* | BKK01760 (BGSC) → SK113 |
| SK130 | *cdaR::kan* | BKK01760 (BGSC) → PY79 |
| SK131 | *cdaR::kan*; *glmR::erm* | SK130 → RB176 |
| SK137 | Δ*cdaA** (markerless) | pDR244 → SK96 |
| SK138 | Δ*cdaA**; *glmR::erm* | BKE34760 (BGSC) → SK137 |
| SK139 | Δ*glmR*; *amyE::*$P_{hyperspank}$*-glmR-T304A spec* | pSK24 → SK35 |
| SK140 | Δ*glmR*; *amyE::*$P_{hyperspank}$*-glmR-T304E spec* | pSK25 → SK35 |
| SK141 | Δ*glmR*; *amyE::*$P_{hyperspank}$*-glmR-R301A spec* | pSK26 → SK35 |
| BLS67 | *glmR::erm* Δ*cdaA** *amyE::*$P_{hyperspank}$*-glmM spec* | pLS29 → SK138 |
| BLS84 | Δ*glmR sacA::*$P_{xyl}$*-glmS* | pLS35 → SK35 |
| BLS95 | Δ*glmR*; *amyE::*$P_{hyperspank}$*-glmR-D38A-D39A spec* | pLS59 → SK35 |
| BLS101 | Δ*glmR*; *amyE::*$P_{hyperspank}$*-glmR-6his spec* | pLS79 → SK35 |
| BLS102 | Δ*glmR*; *amyE::*$P_{hyperspank}$*-glmR-D38A-D39A-6his spec* | pLS80 → SK35 |

**Table S2: Primers used in this study**

| Primer | Sequence |
|---|---|
| oSK16 | AATAA AAGCTT ACATAAGGAGGAACTACT ATGGGACAAAAGCCGAAAATC |
| oSK17 | AATAA GTCGAC TCATTCTTTCAGTAAATCAACAAGAAG |
| oSK44 | AATAA GAATTC CAGAATAAAACAGAGGCGATTTTAGCCTCTG |
| oSK45 | AATAA GCTAGC CATCGATGTCTTCCCCTTTTAATTTCTCATTTTTCAATTAAAACAATAAAACATCC |
| oSK46 | AATAA GCTAGC ATGAGTAAAGGAGAAGAACTTTTC |
| oSK47 | AATAA GGATCC TTATTTGTATAGTTCATCCATGCC |
| oSK48 | AATAAGCTAGCATGGGACAAAAGCCGAAAATC |
| oSK49 | AATAAGGATCCTCATTCTTTCAGTAAATCAACAAGAAG |
| oSK56 | GACGTAATACGTCACGATGCACATAAAGTGGCCTCTC |
| oSK57 | GAGAGGCCACTTTATGTGCATCGTGACGTATTACGTC |
| oSK58 | GACGTAATACGTCACGATGAACATAAAGTGGCCTCTCTTCTTG |
| oSK59 | CAAGAAGAGAGGCCACTTTATGTTCATCGTGACGTATTACGTC |
| oSK60 | CGTATAAAAATGACGTAATAGCTCACGATACACATAAAGTGGCC |
| oSK61 | GGCCACTTTATGTGTATCGTGAGCTATTACGTCATTTTTATACG |
| oSK68 | CAGAGATCAAATTGTAACGTATCAGAATGACGTAATACGTCACGATAC |
| oSK69 | GTATCGTGACGTATTACGTCATTCTGATACGTTACAATTTGATCTCTG |
| oSK70 | CAGAGATCAAATTGTAACGTATAGAAATGACGTAATACGTCACG |
| oSK71 | CGTGACGTATTACGTCATTTCTATACGTTACAATTTGATCTCTG |
| oSK86 | AATAAGTCGACACATAAGGAGGAACTACTATGGGCAAGTATTTTGGAACAG |
| oSK87 | AATAAGCTAGCTTACTCTAATCCCATTTCTGACC |
| oLS69 | AATAATCTAGAACATAAGGAGGAACTACTATGTGTGGAATCGTAGGTTATATCGG |
| oLS70 | AATAACTGCAGTTACTCCACAGTAACACTCTTCGC |
| oLS119 | CCAGAGCTCCCCCCAGCAGCGGCAACTGTTACAA |
| oLS120 | TTGTAACAGTTGCCGCTGCTGGGGGGAGCTCTGG |
| oLS144 | AATAA GTCGAC ACATAAGGAGGAACTACT ATGGGACAAAAGCCGAAAATCGC |
| oLS145 | AATAA GCTAGC TTAGTGATGGTGATGGTGATG TTCTTTCAGTAAATCAACAAGAAGAGAGGCC |

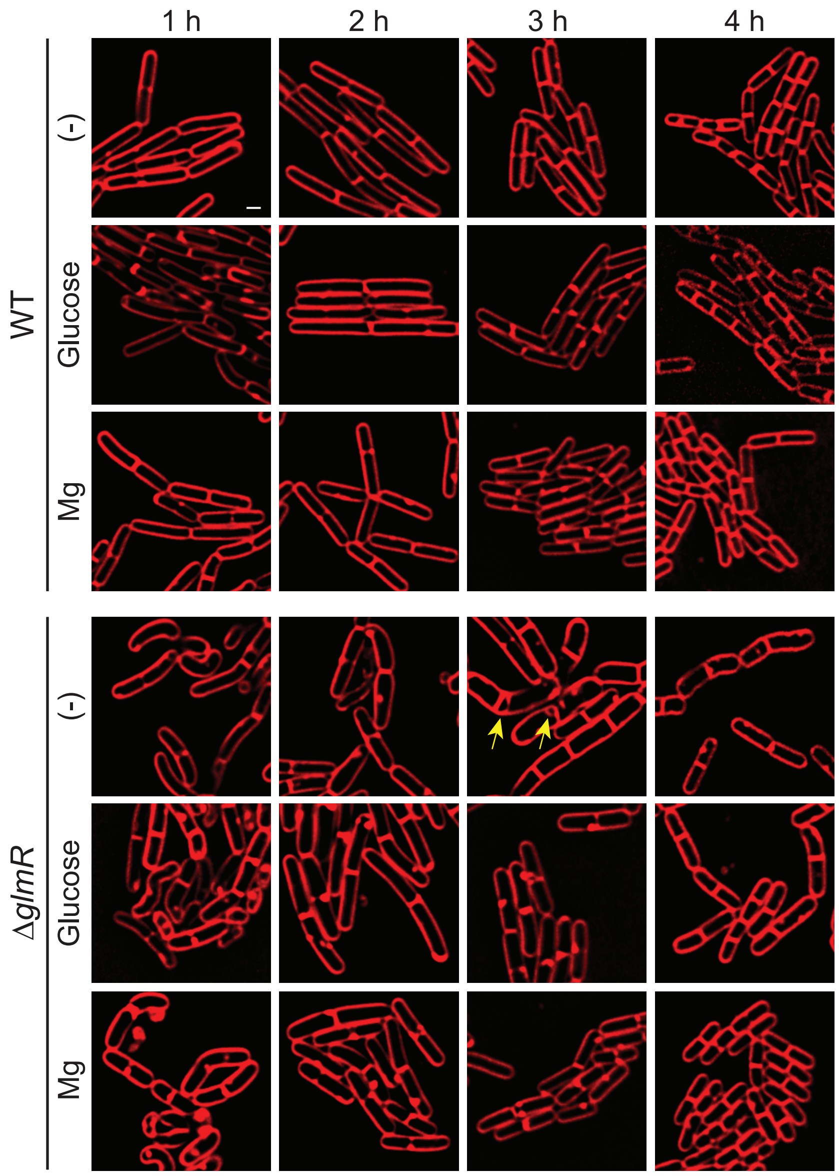


Figure S1

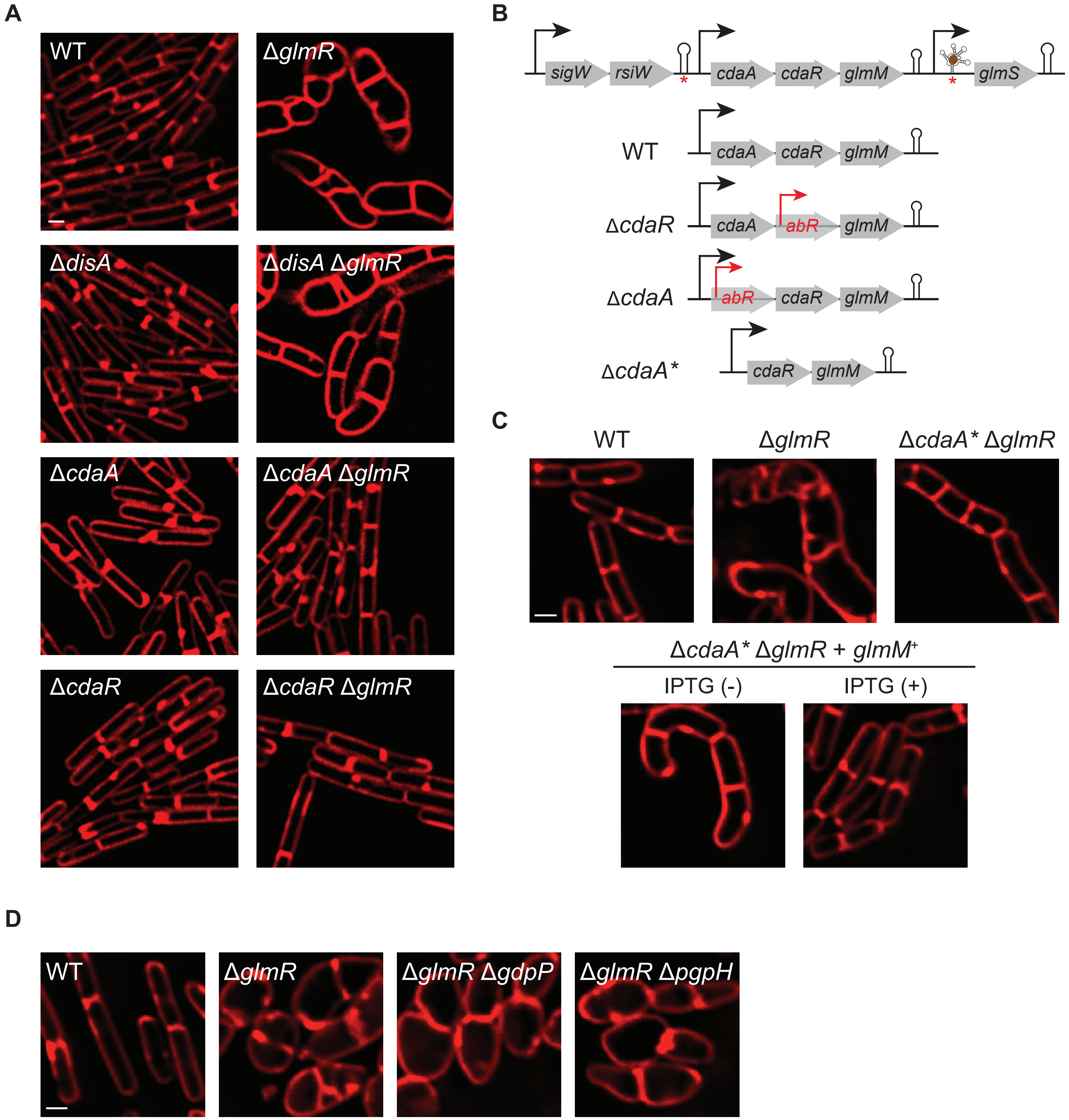
A
WT
ΔglmR
ΔdisA
ΔdisA ΔglmR
ΔcdaA
ΔcdaA ΔglmR
ΔcdaR
ΔcdaR ΔglmR
B
sigW
rsiW
cdaA
cdaR
glmM
glmS
WT
ΔcdaR
abR
ΔcdaA
ΔcdaA*
C
ΔcdaA* ΔglmR
ΔcdaA* ΔglmR + glmM+
IPTG (-)
IPTG (+)
D
ΔglmR ΔgdpP
ΔglmR ΔpgpH

Figure S2

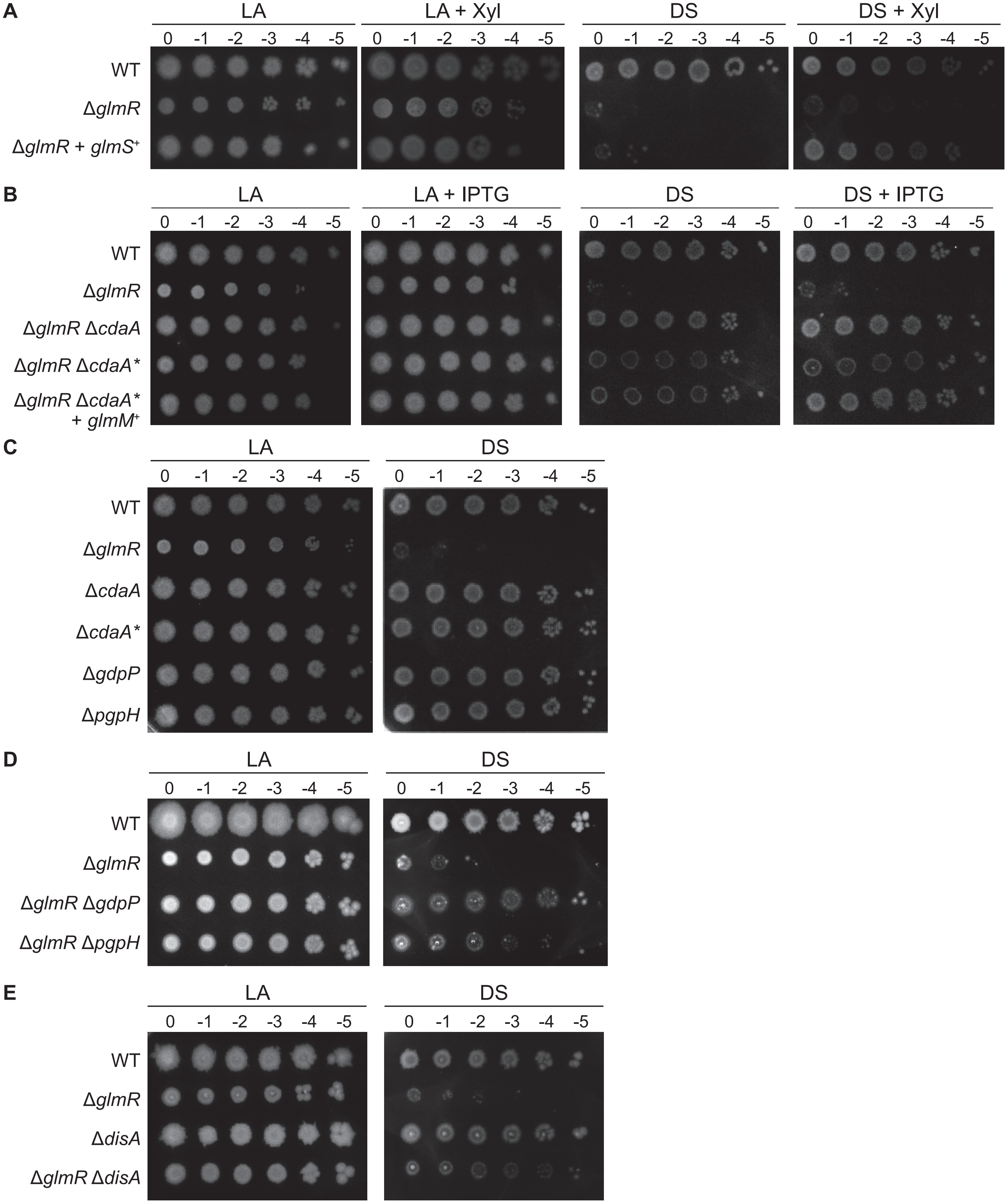
A
LA
LA + Xyl
DS
DS + Xyl
0 -1 -2 -3 -4 -5
WT
ΔglmR
ΔglmR + glmS+
B
LA
LA + IPTG
DS
DS + IPTG
0 -1 -2 -3 -4 -5
WT
ΔglmR
ΔglmR ΔcdaA
ΔglmR ΔcdaA*
ΔglmR ΔcdaA* + glmM+
C
LA
DS
0 -1 -2 -3 -4 -5
WT
ΔglmR
ΔcdaA
ΔcdaA*
ΔgdpP
ΔpgpH
D
LA
DS
0 -1 -2 -3 -4 -5
WT
ΔglmR
ΔglmR ΔgdpP
ΔglmR ΔpgpH
E
LA
DS
0 -1 -2 -3 -4 -5
WT
ΔglmR
ΔdisA
ΔglmR ΔdisA

Figure S3

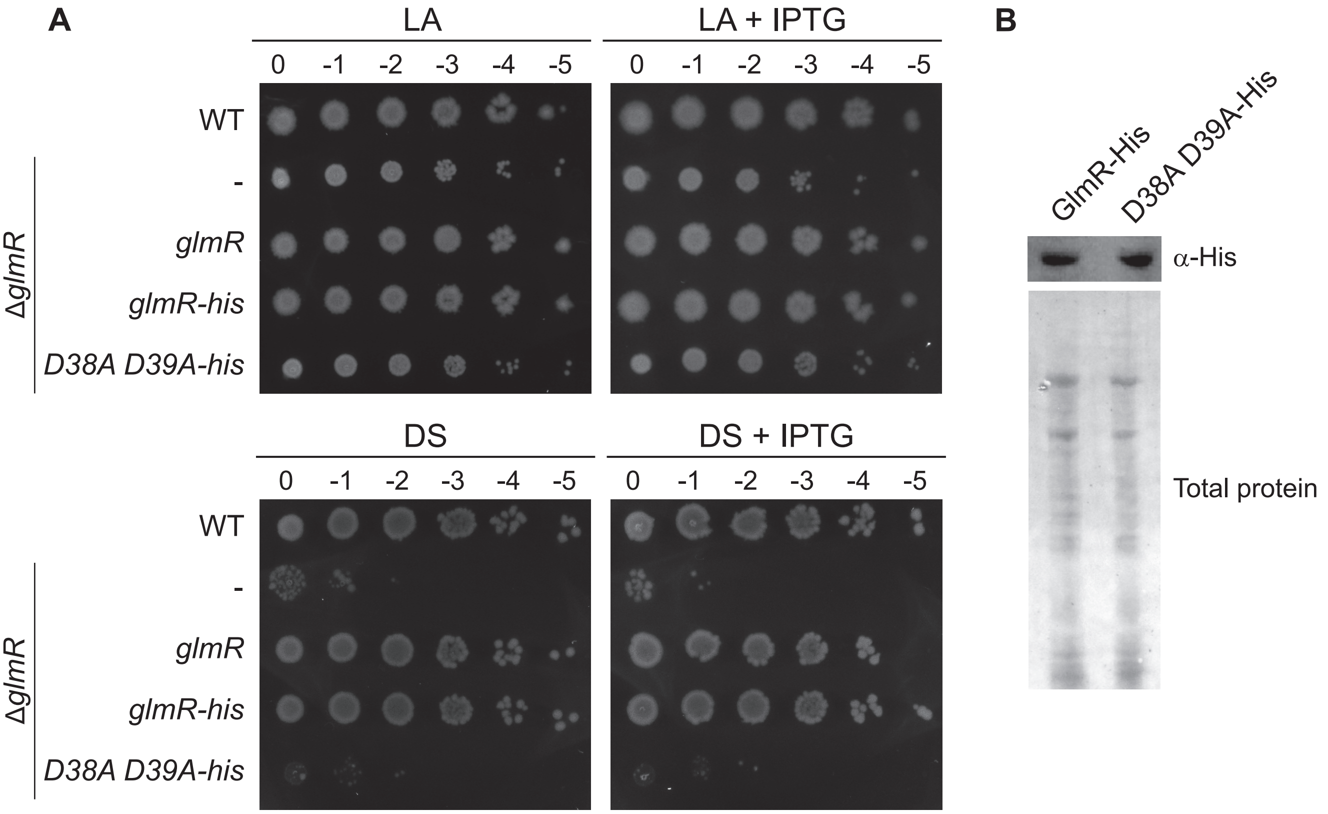
A
LA
LA + IPTG
0 -1 -2 -3 -4 -5
WT
-
glmR
glmR-his
D38A D39A-his
ΔglmR
DS
DS + IPTG
B
GlmR-His
D38A D39A-His
α-His
Total protein

Figure S4

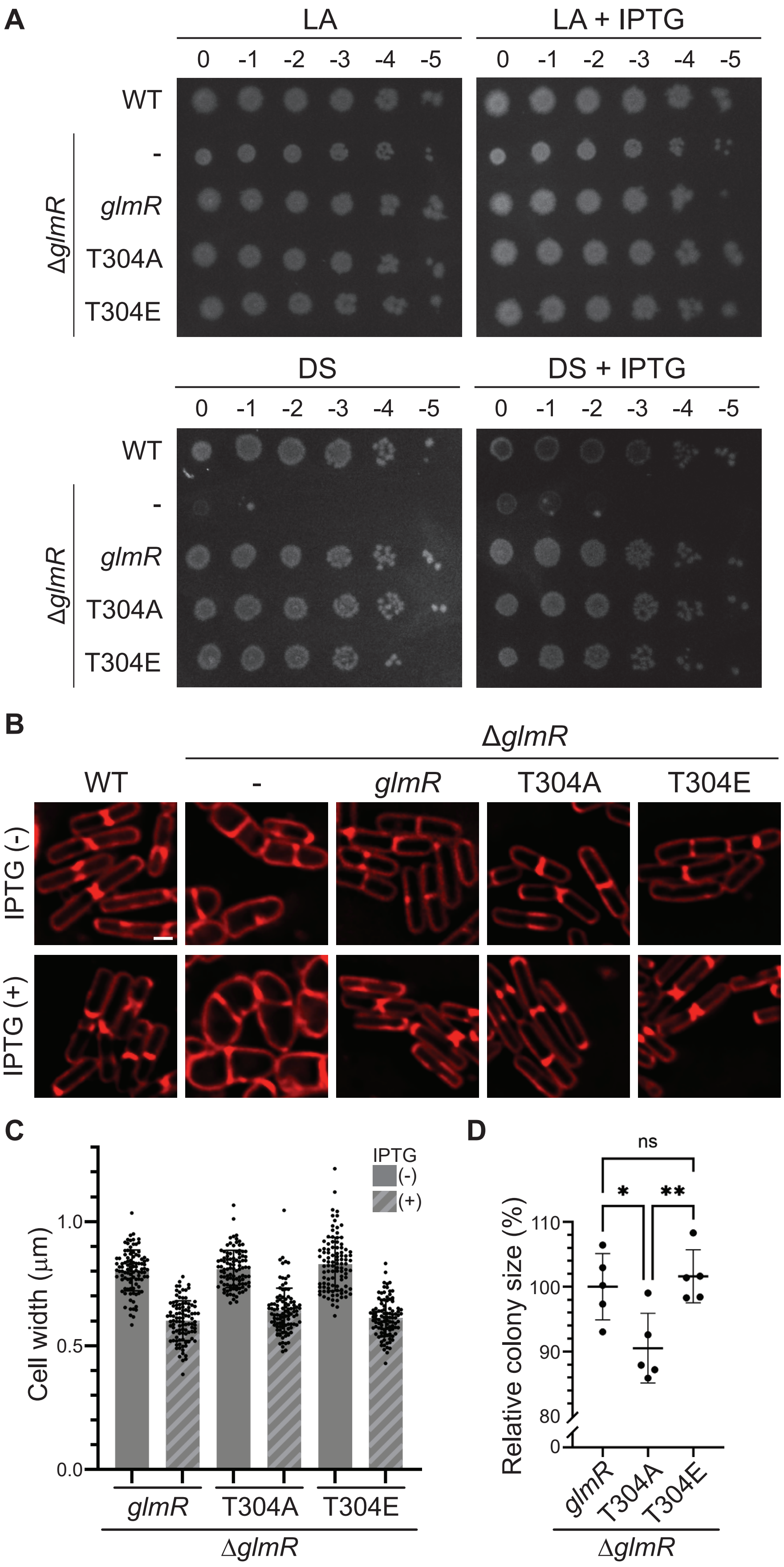

A
LA
LA + IPTG
0 -1 -2 -3 -4 -5
WT
-
glmR
T304A
T304E
ΔglmR
DS
DS + IPTG
B
ΔglmR
WT
-
glmR
T304A
T304E
IPTG (-)
IPTG (+)
C
IPTG
(-)
(+)
Cell width (μm)
1.0
0.5
0.0
glmR
T304A
T304E
ΔglmR
D
ns
*
**
Relative colony size (%)
110
100
90
80
0
glmR
T304A
T304E
ΔglmR


Figure S5

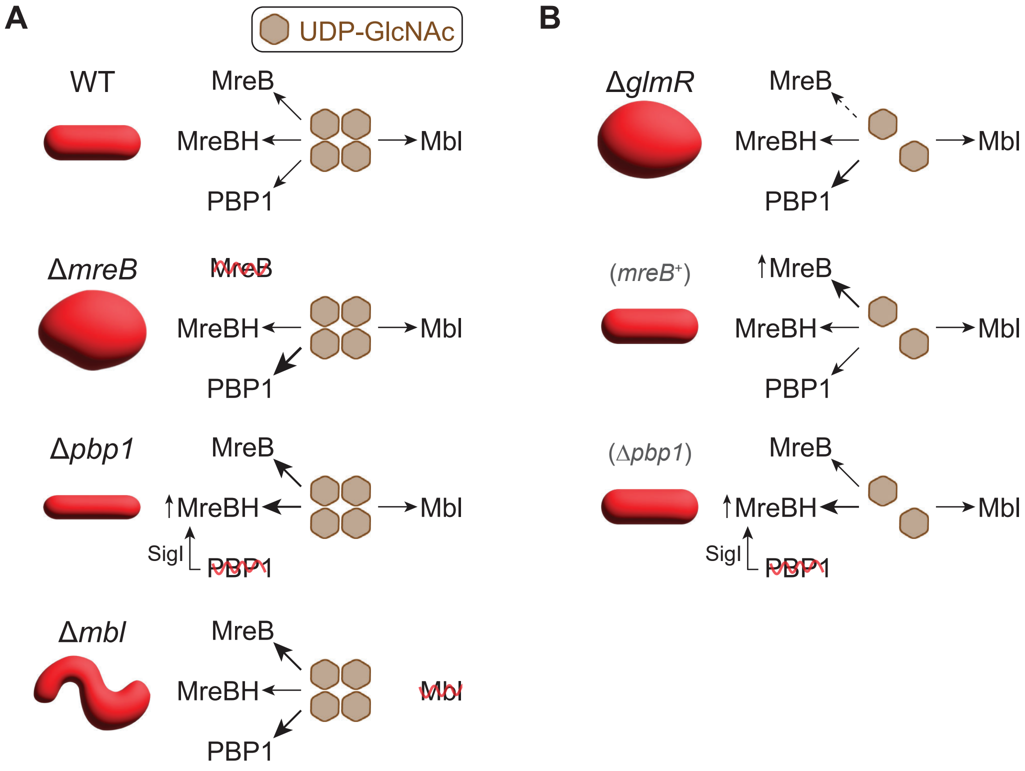
A
UDP-GlcNAc
WT
MreB
MreBH
Mbl
PBP1
ΔmreB
MreB
MreBH
Mbl
PBP1
Δpbp1
MreB
MreBH
Mbl
SigI
PBP1
Δmbl
MreB
MreBH
Mbl
PBP1
B
ΔglmR
MreB
MreBH
Mbl
PBP1
(mreB+)
MreB
MreBH
Mbl
PBP1
(Δpbp1)
MreB
MreBH
Mbl
SigI
PBP1

Figure S6
